# Molecular Logic of Synaptic Diversity Between *Drosophila* Tonic and Phasic Motoneurons

**DOI:** 10.1101/2023.01.17.524447

**Authors:** Suresh K. Jetti, Andrés B. Crane, Yulia Akbergenova, Nicole A. Aponte-Santiago, Karen L. Cunningham, Charles A. Whittaker, J. Troy Littleton

## Abstract

Although neuronal subtypes display unique synaptic organization and function, the underlying transcriptional differences that establish these features is poorly understood. To identify molecular pathways that contribute to synaptic diversity, single neuron PatchSeq RNA profiling was performed on *Drosophila* tonic and phasic glutamatergic motoneurons. Tonic motoneurons form weaker facilitating synapses onto single muscles, while phasic motoneurons form stronger depressing synapses onto multiple muscles. Super-resolution microscopy and *in vivo* imaging demonstrated synaptic active zones in phasic motoneurons are more compact and display enhanced Ca^2+^ influx compared to their tonic counterparts. Genetic analysis identified unique synaptic properties that mapped onto gene expression differences for several cellular pathways, including distinct signaling ligands, post-translational modifications and intracellular Ca^2+^ buffers. These findings provide insights into how unique transcriptomes drive functional and morphological differences between neuronal subtypes.

## Introduction

Although molecular pathways driving neuronal development and function are well characterized, how diversity in cell subtypes across the brain is established and maintained is poorly understood. With the advent of single-cell transcriptomics, mRNA expression profiles that contribute to distinct neuronal morphology, connectivity and function are being identified (Belgard et al., 2011; Brunet Avalos et al., 2019; Corrales et al., 2022; Darmanis et al., 2015; Fuzik et al., 2016; Lake et al., 2016; Poulin et al., 2016). One central feature of diversity is the unique synaptic properties observed across neuronal classes. Synapses represent fundamental building blocks of information processing, exhibiting distinct synaptic vesicle (SV) release probability (*Pr*), response kinetics and short-term plasticity (Dittman et al., 2000; Fioravante and Regehr, 2011; Grande and Wang, 2011; Millar et al., 2002). Notable examples of synaptic diversity include *C. elegans* AWC and ASH olfactory neurons (Ventimiglia and Bargmann, 2017), invertebrate tonic and phasic motoneurons (MNs) (Atwood, 2008; Lnenicka and Keshishian, 2000; Takeda and Kennedy, 1964), zebrafish ON and OFF bipolar cells (Odermatt et al., 2012), and mammalian hippocampal, cerebellar and central auditory neurons (Dittman et al., 2000; Regehr, 2012; Rollenhagen and Lübke, 2006). Functional heterogeneity in synaptic transmission contributes to temporal coding and frequency filtering, processing of multisensory information, and various circuit computations (Abbott and Regehr, 2004; Chabrol et al., 2015; Chadderton et al., 2014; Díaz-Quesada et al., 2014). Despite these roles, the differentially expressed genes (DEGs) that collectively specify differences in synaptic structure and output are largely unknown.

Neurons displaying tonic or phasic patterns of synaptic output are conserved from invertebrates to mammals and often co-innervate postsynaptic targets (Aponte-Santiago and Littleton, 2020; Atwood and Karunanithi, 2002; Ellwood et al., 2017; Ventimiglia and Bargmann, 2017). Tonic and phasic synapses can increase robustness of information processing by acting as high and low-pass filters, respectively (Abbott and Regehr, 2004; Jackman and Regehr, 2017). *Drosophila* larvae contain MNs with either tonic or phasic output that co-innervate muscles to form glutamatergic neuromuscular junctions (NMJs) with unique synaptic features (Aponte-Santiago and Littleton, 2020; Lnenicka and Keshishian, 2000). Tonic MNs have bigger “type Ib” boutons and typically innervate single muscles to act as primary drivers of contraction (Hoang and Chiba, 2001; Johansen et al., 1989; Newman et al., 2017). Phasic MNs form smaller “type Is” boutons that innervate and coordinate contraction of muscle subgroups. Phasic MNs form active zones (AZs) that display higher *Pr* and show synaptic depression during high frequency stimulation (Aponte-Santiago et al., 2020; Genç and Davis, 2019; Karunanithi et al., 2020; Lu et al., 2016; Newman et al., 2017). In contrast, tonic MNs display lower synaptic output and undergo facilitation. Tonic and phasic MNs also show differences in morphology, presynaptic inputs, membrane excitability and synaptic plasticity (Aponte-Santiago and Littleton, 2020; Choi et al., 2004; Clark et al., 2018; Couton et al., 2015; Lnenicka, 2020; Zarin et al., 2019). The molecular logic that governs these differences is largely unknown, providing a system to characterize DEGs that establish and maintain these distinct properties.

In this study we combined single-neuron PatchSeq RNA profiling with quantal imaging, stimulated emission depletion (STED) nanoscopy, transmission electron microscopy (TEM), optogenetics, electrophysiology, and genetic manipulations to identify and dissect molecular components contributing to structural and functional diversity of *Drosophila* tonic and phasic MNs. RNAseq identified distinct transcriptional profiles for each MN subtype, while genetic analyses indicate multiple DEGs contribute to functional and morphological differences in AZ organization, *Pr*, Ca^2+^ buffering and reliance on AZ scaffolds.

## Results

### Tonic and phasic MNs exhibit diversity in morphology and synaptic properties

The *Drosophila* larval motor system contains glutamatergic (tonic type Ib and phasic type Is) and neuromodulatory (type II and III) MNs that control locomotion. In each abdominal hemisegment, ∼30 Ib MNs individually innervate the 30 body wall muscles, while two Is MNs innervate the dorsal or ventral muscles. Ib and Is MNs display differences in excitability, morphology and synaptic strength as previously described (reviewed in Aponte-Santiago and Littleton, 2020). We quantified their structural and functional synaptic properties as a foundation for characterizing DEGs that contribute to these features. We focused on tonic MN1-Ib (aCC) that innervates muscle 1 and phasic MNISN-Is (RP2, hereafter referred to as Is) that innervates dorsal muscles 1, 2, 3, 4, 9, 10, 11, 19 and 20 (Figure 1A). Gal4 drivers for these neurons are available and facilitate cell-type specific manipulations (Supplemental Figure 1A, Aponte-Santiago et al., 2020; Pérez-Moreno and O’Kane, 2019). MN1-Ib and MN4-Ib have similar properties, and comparisons of MN4-Ib with Is were performed in some experiments as segregation of their synaptic arbors at muscle 4 facilitated imaging studies.

**Figure 1.**
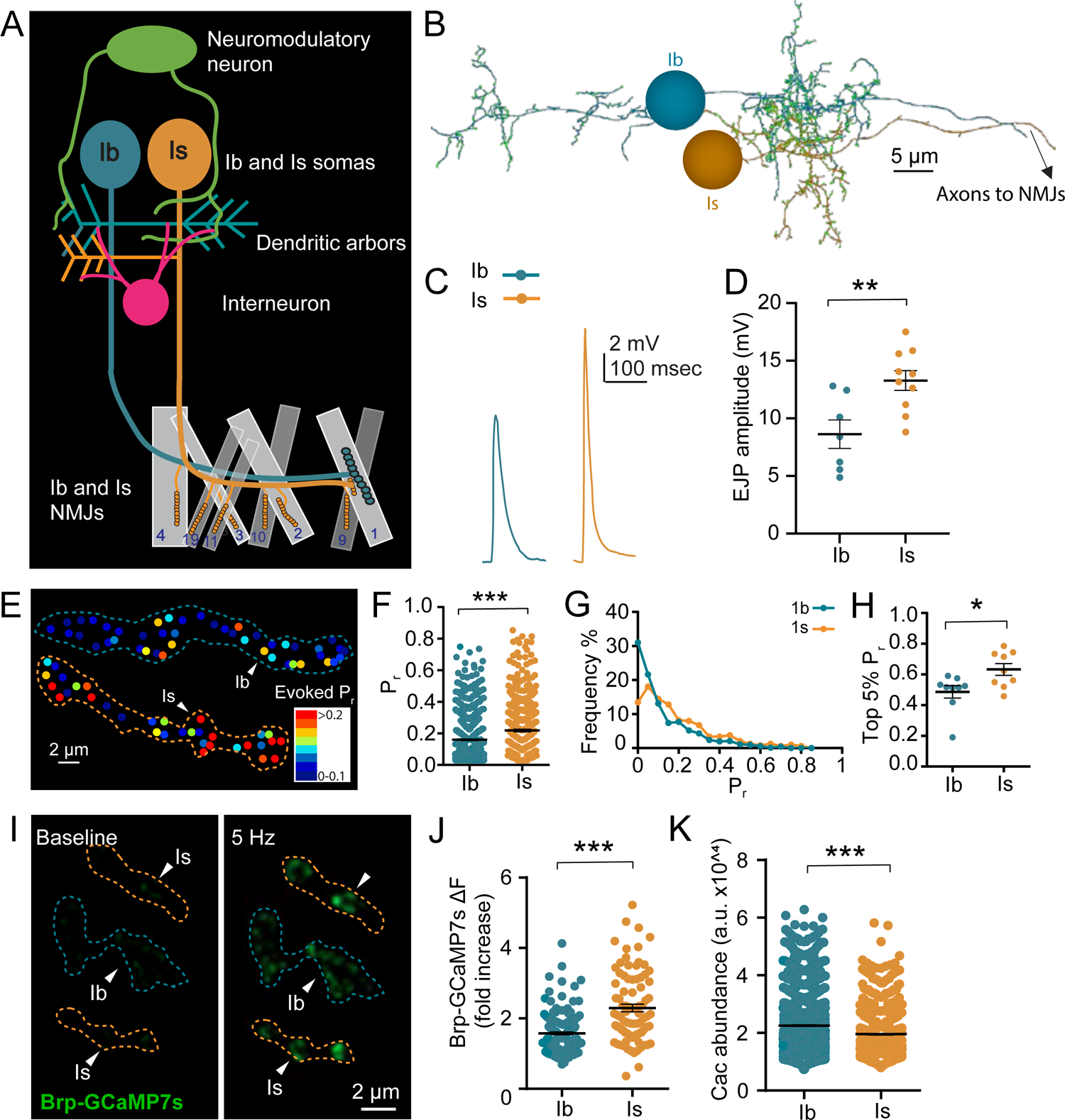
Synaptic differences in tonic Ib and phasic Is motoneurons. (A) Model of larval MN1-Ib and MNISN-Is. Somas are located in the VNC, with the Ib exclusively innervating muscle 1, while the Is innervates the dorsal muscle subgroup. (B) EM reconstruction of MN1-Ib and MNISN-Is from 1^st^ instar abdominal segment A1. Synaptic inputs onto Ib and Is dendrites within the VNC are noted by green circles. Ib forms both ipsilateral and contralateral dendritic branches that cross the midline, while the Is dendritic arbor resides in the ipsilateral VNC. (C) Representative EJP traces at 3^rd^ instar muscle 1 evoked by optogenetic stimulation of UAS-Channelrhodopsin driven with MN1-Ib or Is Gal4 drivers. (D) Quantification of muscle 1 EJP amplitude for optical stimulation of Ib versus Is MNs (n=7 larvae). (E) Representative evoked AZ *Pr* heatmap at muscle 4 following quantal imaging of postsynaptic during 0.3 Hz stimulation. (F) Quantification of *Pr* for Ib and Is AZs at muscle 4 NMJs from quantal imaging (Ib: 716 AZs from 9 NMJs, 4 larvae; Is: 476 AZs from 9 NMJs, 4 larvae). (G) Histogram of single AZ *Pr* for Ib and Is NMJs at muscle 4. (H) Quantification of average *Pr* for the strongest 5% of Ib or Is AZs at individual muscle 4 NMJs (n=9 NMJs, 4 larvae). (I) Representative images of resting (left) and 5 Hz evoked (right) Ca^2+^ signals at Ib and Is NMJs at muscle 4 in larvae expressing Brp-GCaMP7s. (J) Quantification of fold-increase in Ca^2+^ levels during 5 Hz stimulation of Ib and Is NMJs at muscle 4 in larvae expressing Brp-GCaMP7s (Ib: n=169 AZs from 7 NMJs, 3 larvae; Is: n=87 AZs from 6 NMJs, 3 larvae). (K) Quantification for Cac abundance (in a.u) at Ib and Is AZs at muscle 4 (Ib: n=1600 AZs from 12 NMJs, 4 larvae; Is: n=743 AZs from 12 NMJs, 4 larvae). Data reports mean ± SEM. Student t-test, \**p* < 0.05, \*\**p* < 0.01, \*\*\**p* < 0.001, ns not significant.

To quantify MN1-Ib and Is terminal morphology, synaptic NMJ area and AZ number were assayed at 3^rd^ instar NMJs (Supplemental Figure 1B-E). MN1-Ib had bigger boutons and a 4.4-fold larger NMJ field with 3.8-fold more AZs compared to Is at muscle 1 (p<0.0001). However, when all postsynaptic muscles innervated by the dorsal Is were included, the total synaptic area of the Is MN was 4.6-fold larger with 4.2-fold more AZs (p<0.0001). To examine dendritic inputs, the annotated EM connectome of abdominal segment one in the 1^st^ instar larval ventral nerve cord (VNC) was analyzed (Milyaev et al., 2012; Ohyama et al., 2015; Schneider-Mizell et al., 2016; Zarin et al., 2019). The dendrites of MN1-Ib bifurcate into ipsilateral and contralateral branches and contain 201 input synapses from 31 upstream neurons, while the Is dendritic arbor is confined to the ipsilateral side of the VNC with 137 input synapses from 21 neurons at this developmental stage (Figure 1B). MN1-Ib and Is receive inputs from both shared and distinct pre-motor and interneuron pools that drive their sequential recruitment during locomotion (Fushiki et al., 2016; Heckscher et al., 2015; Schneider-Mizell et al., 2016; Zwart et al., 2016). In summary, MN1-Ib has ∼75% fewer output synapses and ∼50% more input synapses than the dorsal Is, indicating the transcriptomes of each MN subtype must support distinct pre- and post-synaptic development and maintenance requirements.

To assay presynaptic output, UAS-Channelrhodopsin-2 was expressed with MN1-Ib or Is specific Gal4 drivers and wide-field optogenetic NMJ stimulation was performed. Optically evoked excitatory junction potentials (EJPs) at muscle 1 were 54% larger following Is stimulation compared to Ib (p<0.01, Figure 1C, D), consistent with prior studies (Genç and Davis, 2019). Quantal imaging was performed using myristoylated-GCaMP7s to examine synaptic output at individual AZs. Expression of this genetically encoded Ca^2+^ indicator in muscle 4 allowed detection of single SV fusion events at defined AZs (Akbergenova et al., 2018; Melom et al., 2013). Similar to recent studies (Newman et al., 2017; Sauvola et al., 2021), Is AZs showed a 1.6-fold higher *Pr* than Ib AZs (p<0.0001, Figure 1E-G). Is MNs also had a smaller population of functionally silent AZs (Is, 5.6%; Ib, 17.1%, p<0.001) and a higher *Pr* ceiling for the strongest release sites (p<0.02, Figure 1H). To assay presynaptic Ca^2+^ influx at Ib and Is AZs, MNs were stimulated at 5 Hz in transgenic larvae expressing GCaMP7s fused to the N-terminus of the AZ protein Bruchpilot (Brp), which resides near presynaptic voltage-gated Ca^2+^ channels (VGCCs) encoded by *Cacophony* (Cac) (Kittel et al., 2006). Is AZs displayed 46% more presynaptic Ca^2+^ influx than Ib (p<0.0001, Figure 1I, J). To determine if elevated Ca^2+^ influx was secondary to more Ca^2+^ channels, quantification of CRISPR-tagged Cac-GFP AZ abundance was performed as previously described (Akbergenova et al., 2018; Cunningham et al., 2022). A 13% reduction in Cac channels was observed at Is AZs (p<0.0001, Figure 1K), indicating Is AZs display greater Ca^2+^ influx and higher *Pr* despite having fewer Cac channels.

### The nanoscopic organization of Ib and Is AZs are distinct

*Drosophila* AZs form a mushroom-shaped structure composed of Brp and other scaffolding proteins that appear as an electron dense T-bar in TEM and a ring-like donut after STED imaging of Brp (Fouquet et al., 2009). To identify structural differences between Is and Ib AZs, Brp immunostaining and STED imaging were performed. Unlike the larger donut-shaped AZs observed in Ib, Is AZs displayed a triangular appearance (Figure 2A) with reduced circularity index (CI) (p<0.0001, Figure 2B) and 37% smaller area (p<0.0001, Figure 2C), suggesting a distinct nanoscopic architecture. To examine Ca^2+^ channel distribution at the AZ center, dual-color STED imaging was performed for Brp and GFP-tagged endogenous Cac (Gratz et al., 2019). Cac channel distribution was not significantly different (Figure 2D, E) and increased Brp area correlated with higher Cac area for both Ib and Is AZs (Figure 2F).

**Figure 2.**
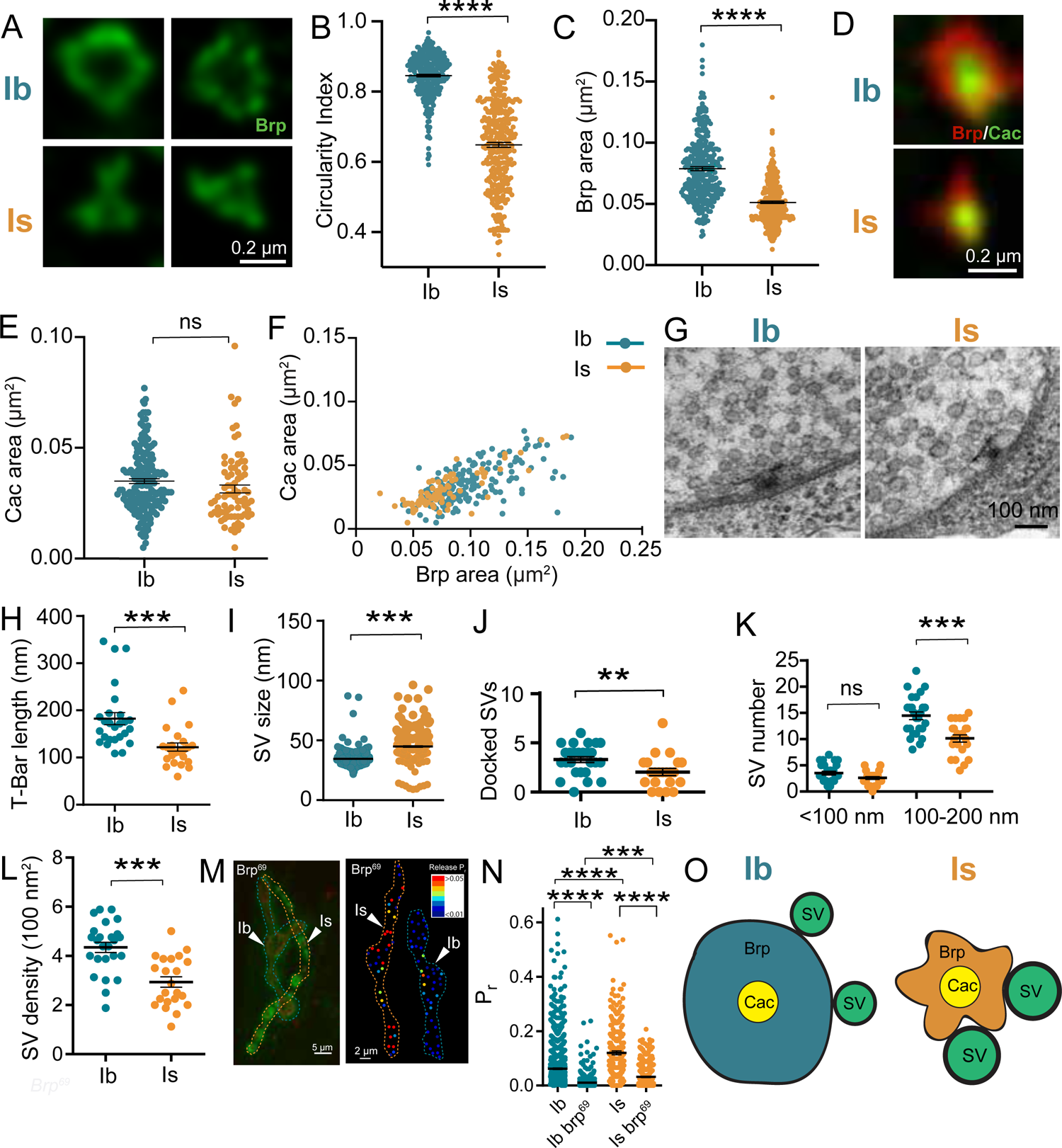
AZ nanoscopic diversity between Ib and Is AZs. (A) Representative STED images showing Ib (upper panel) and Is (lower panel) AZs immunolabeled with anti-Brp. (B) Quantification of circularity index differences between Ib and Is AZs using STED. (C) Quantification of Brp area differences between Ib and Is AZ using STED (n=314 Ib AZs from 7 larva and 345 Is AZs, 9 larva for panels B and C). (D) Representative STED images of endogenous CRISPR GFP-tagged Cac channels and Brp at Ib and Is AZs. (E) Quantification of Cac area at Ib and Is AZs using STED. (F) Quantification of Cac area vs. Brp area at Ib and Is AZs using STED (n=175 Ib AZs from and 79 Is AZs, 4 larva for panels E and F). (G) Representative TEM images of Ib and Is T-bar organization. (H) Quantification of T-bar length at Ib and Is AZs (Ib: n=26 micrographs; Is: n=24 micrographs). (I) Quantification of SV diameter at Ib and Is synapses (Ib: n=298 Ib SVs from 10 micrographs; Is: n=338 Is SVs from 10 micrographs). (J) Quantification of docked SVs at Ib and Is AZs (Ib: n=26 micrographs; Is: n=22 micrographs). (K) Quantification of SV distribution around Ib and Is T-bars (Ib: n=26 micrographs; Is: n=22 micrographs). (L) Quantification of SV density at Ib and Is boutons (Ib: n=25 micrographs; Is: n=22 micrographs). (M) Representative image (left) and evoked *Pr* map (right) of quantal imaging with myristoylated-GCaMP7s at Ib and Is AZs in *brp*^69^ mutants. (N) Quantification for evoked *Pr* in control and *brp*^69^ mutants (Con Ib: n=1047 AZs from 7 NMJs, 6 larvae; *brp*^69^ Ib: n=502 AZs from 6 NMJs, 6 larvae; Con Is: n=222 AZs from 6 NMJs, 6 larvae; *brp*^69^ Is: n=301 AZs from 6 NMJs, 6 larvae). Data reports mean ± SEM. Student t-test, \**p* < 0.05, \*\**p* < 0.01, \*\*\**p* < 0.001, n.s not significant. (O) Model summarizing nanoscopic differences in Ib and Is AZ organization. Is AZs have bigger SVs, enhanced Ca^2+^ influx and a smaller Brp cytomatrix compared to the larger Ib AZs. Data represents mean ± SEM. One-way ANOVA with Tukey correction was used to determine significance in K and N panels. Student t-test was used for two bar graphs. \**p* < 0.05, \*\**p* < 0.01, \*\*\**p* < 0.001, \*\*\*\**p* < 0.0001, ns not significant.

Given the role of Brp in recruiting SVs to AZs (Matkovic et al., 2013), the distinct Brp AZ architecture detected by STED might alter the number or distribution of SVs at release sites that could contribute to differences in synaptic strength. To quantify SV distribution and AZ ultrastructure, TEM analysis of 3^rd^ instar larval Ib and Is synaptic boutons was performed (Figure 2G). Consistent with the smaller Brp area, T-bar length was 33% shorter at Is AZs (p<0.001, Figure 2H). As previously described (Karunanithi et al., 2002), SVs in Is boutons were 30% larger than those at Ib synapses (p<0.0001, Figure 2I). In spite of higher *Pr*, Is AZs contained fewer docked SVs within 40 nm of the electron dense synaptic cleft (p<0.001, Figure 2J) and less SVs clustered within 100-200 nm of the T-bar (p<0.0001, Figure 2K). In addition, Ib MNs had a 2-fold increase in overall SV density within synaptic boutons (p<0.01, Figure 2L), suggesting greater potential for sustained release during stimulation to support their tonic output. If the distinct morphology of Ib and Is AZs observed by STED and TEM contributes to *Pr* differences, the two MN subtypes might display differential sensitivity to Brp disruption. To test Brp-dependent SV release, *Pr* mapping was performed in *brp^69^* null mutants at Ib and Is NMJs. Although evoked release was reduced at both terminals in *brp^69^*, Is AZs were slightly more resistant and maintained higher *Pr* than Ib AZs (p<0.001, Figure 2M, N). These data suggest baseline *Pr* is Brp-dependent in both MN subtypes, with cell type-specific *Pr* differences requiring factors in addition to Brp-mediated structural changes. Together, STED and TEM analysis indicate the nanoscale organization of the AZ cytomatrix and associated SV pools differs at Ib and Is AZs. Is AZs display higher *Pr* despite fewer docked SVs, suggesting a model where individual SV fusogenicity and greater Ca^2+^ influx represent key drivers for enhanced release (Figure 2O).

### PatchSeq RNA profiling identifies distinct transcriptional profiles for Ib and Is MNs

To examine pathways that establish these distinct synaptic properties, the transcriptomes of the two MN subtypes were compared. MN1-Ib and Is are among the first MNs born during embryogenesis and arise from distinct segmental neuroblast lineages. MN1-Ib, along with several interneurons and glia, derive from neuroblast NB1-1, while the Is and a few additional MNs and interneurons are descendants of neuroblast NB4-2. The initial transcriptional wave for synaptic genes occurs during embryogenesis from 12-24 hours as the nervous system forms (Gerstein et al., 2014). NMJ growth and AZ addition increase dramatically during the 2^nd^ and 3^rd^ instar stages (Akbergenova et al., 2018; Zito et al., 1999), and MNs continue to express genes required for maintaining their synaptic properties and forming new AZs (Cunningham et al., 2022). Our synaptic studies were performed in 3^rd^ instar larvae that are ∼5 days beyond embryogenesis, and we focused on characterizing Ib and Is transcriptomes during this developmental period.

Single neuron PatchSeq RNA profiling was performed using a method modified from adult *Drosophila* neurons (Crocker et al., 2016). MN-Ib and Is cells were labeled with GFP using specific Gal4 drivers to identify their cell bodies on the ventral surface of the 3^rd^ instar VNC (Aponte-Santiago et al., 2020). RNA profiles for two postsynaptic partners, muscle 1 (innervated by MN1-Ib and Is) and muscle 4 (innervated by MN4-Ib and Is), were also determined. Cytosolic and nuclear content from individual cell bodies of 101 Is, 105 MN-Ib and 35 muscles was removed using whole-cell electrodes (Figure 3A) and high-resolution paired-end deep single-cell RNA sequencing using Smart Seq-2 was performed (Trombetta et al., 2014). We termed this approach ‘Isoform-PatchSeq’ as the method resulted in ∼4 million reads per cell and allowed identification of DEG profiles and splice isoform variants at single cell resolution. Read counts for all genes and isoforms, along with a regularized logarithm transformation (rlog) in DeSeq2, is shown in Supplemental Table 1 (Love et al., 2014). Unsupervised hierarchical clustering and tSNE analyses indicated the transcriptomic profiles of MN-Ib, Is, and muscles clustered into separate groups (Figure 3B). Comparison of the two MN populations identified 422 genes upregulated in Ib MNs and 402 genes upregulated in Is MNs with an adjusted p-value of <0.05 (Figure 3C). A larger set of ∼5,000 DEGs were identified between MNs and muscles.

**Figure 3.**
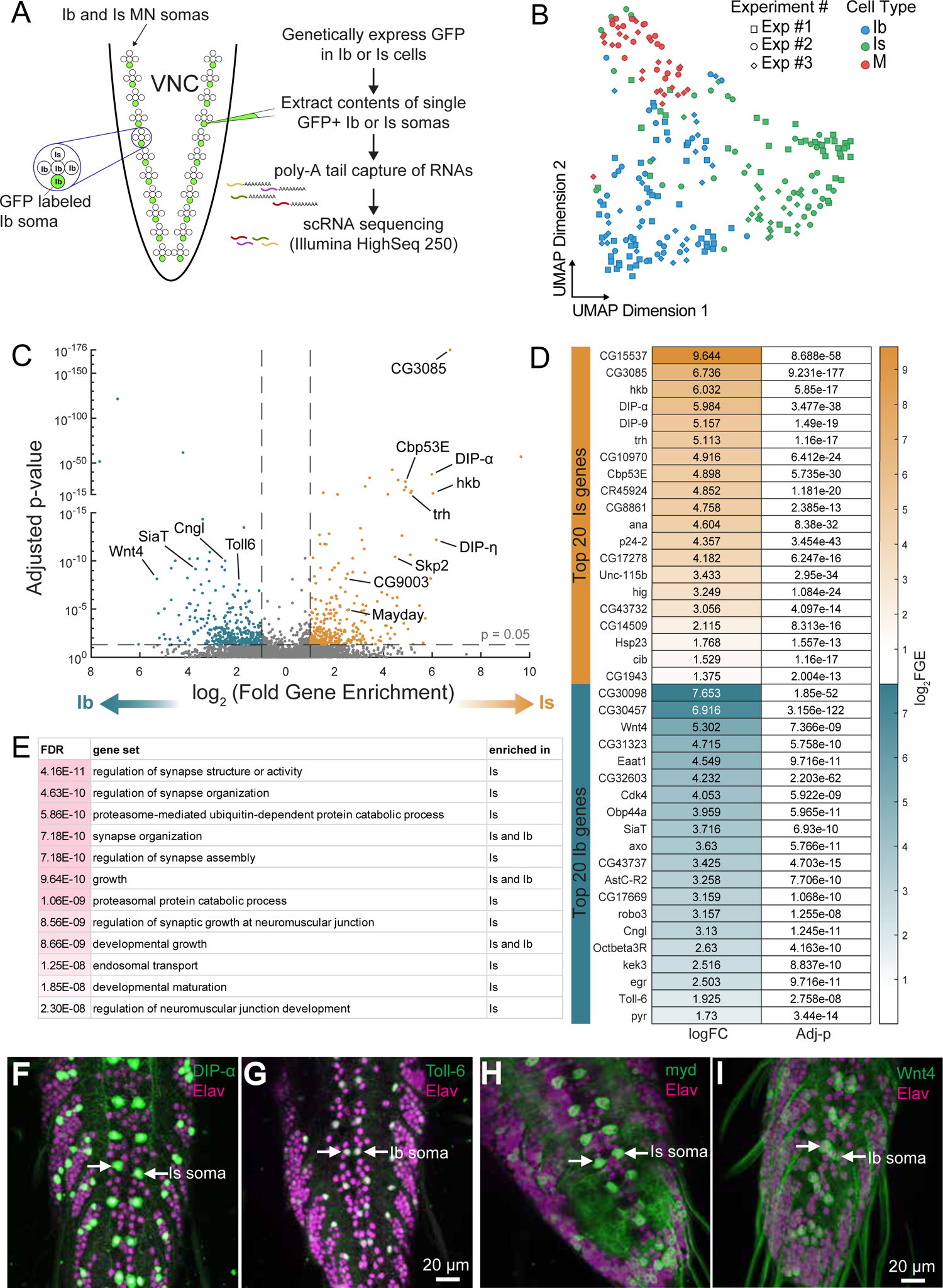
Isoform-PatchSeq analysis of Ib and Is motoneurons. (A) Scheme for single MN Isoform PatchSeq approach. (B) Visualization of Ib and Is MNs compared to larval muscles 1 and 4 using t-distributed stochastic neighbor embedding (tSNE). Each dot represents a single MN or individual muscle coded in blue (Ib), green (Is) or red (muscle). Distinct experiments and sequencing runs are denoted by symbol shape (n=101 Is MNs, 105 MN-Ibs and 35 muscle samples). (C) Volcano plot of gene expression differences between MN1-Ib and Is, with each dot representing a single gene. Log2 change (FC) and adjusted p-values were plotted on the x and y axes, respectively. Genes upregulated in Ib and Is are color-coded in turquoise and orange. (D) Heat map of the top 20 DEGs in MN1-Ib and Is. (E) Top gene ontology (GO) categories for DEGs enriched in Is or Ib MNs. (F) Representative images of anti-GFP staining of larval VNCs in protein-trap lines for Is-enriched DIP-α and Ib-enriched Toll-6. Elav immunostaining to label all neurons is shown in magenta. (G) Representative images of anti-GFP staining of larval VNCs in Trojan-Gal4 lines for Is-enriched Mayday (myd) and Ib-enriched Wnt4. Elav immunostaining to label all neurons is shown in magenta.

Isoform-PatchSeq identified 11 previously reported Ib and Is DEGs (*Proc, hkb, trh, vvl, DIP-α, zfh1, SiaT, Toll-6, pav, grn and dpr4*), in addition to many unknown DEGs. The 20 most statistically significant upregulated DEGs in Is MNs included those encoding the Ig domain containing axonal pathfinding proteins DIP-α and DIP-θ, the transcription factors Huckebein (hkb) and Trachealess (trh), the Ca^2+^ binding protein Calbindin 53E (Cbp53), the secreted glycoprotein Anachronism (ana) and serine protease Pegasus (CG17278), the synaptic cleft protein Hig, the intracellular transport protein p24-2, the actin-binding proteins Cib and Unc-115b, the microtubule binding protein JPT1 (CG1943), the neuronal excitability regulator Julius Seizure (CG14509), the chaperone Hsp23 and the long non-coding RNA CR45924 (Figure 3D). Among the top 20 upregulated DEGs in Ib MNs were those encoding the cell surface proteins Toll-6, Kek-3, Robo3 and Axotactin (axo), the signaling ligands Wnt4, Pyramus (FGF ligand) and Eiger (TNF ligand), the glutamate transporter Eaat1, the cyclin Cdk4, the dynein assembly factor Dnaaf3 (CG43737), the sialyltransferase ST6Gal (SiaT), the neuropeptide receptors AstC-R2 and Oct*β*3R, the cyclic nucleotide-gated ion channel CNGL, and the long non-coding RNA CR45139. CG3085 (encoding a tektin family cytoskeletal protein) and CG15537 (homologous to corticotropin-releasing hormone binding protein) were Is MN-specific, while CG30457 (unknown function) and CG30098 (endopeptidase) were Ib MN-specific, showing binary ON/OFF expression between Is and Ib MNs. Isoform Patch-Seq also identified the major splice isoforms for every gene expressed in MNs, as well as splicing differences between MN1-Ib and Is (Supplemental Table 2, Supplemental Item 1). Although functional analysis of splicing differences will require further study, the glutamate receptor accessory protein Neto displayed differential isoform expression, with Neto-*α* being specifically upregulated in MN1-Ib. mRNAs encoding splice isoforms for two well-characterized AZ proteins, Brp^short^/Brp^long^ and Unc13A/Unc13B, were not different between Ib and Is. Gene Ontology (GO) analysis for all DEGs indicated synapse structure and function, along with proteasomal processing and developmental growth, were the most significant categories differentiating MN1-Ib and Is (Figure 3E), consistent with roles in establishing or maintaining their unique structural and functional features.

To analyze protein expression for a subset of DEGs, immunofluorescence experiments were performed using GFP protein trap lines (Nagarkar-Jaiswal et al., 2015). Prior studies found the Ig superfamily protein DIP-α labels Is MNs (Aponte-Santiago et al., 2020; Ashley et al., 2019), while the neurotrophin receptor Toll-6 marks 1b MNs (McIlroy et al., 2013). Isoform-PatchSeq indicated *DIP-α* was enriched >63-fold in Is MNs and *Toll-6* was enriched ∼4-fold in Ib MNs. GFP trap lines confirmed cell-type specific protein expression from these genes (Figure 3F). Trojan-Gal4 lines inserted within newly discovered DEGs also showed specific expression in Ib versus Is cells as expected (Figure 3G, Supplemental Figure 2). *Wnt4* (∼40-fold RNA increase in Ib) and *DIP-γ* (>3-fold RNA increase in Ib) were preferentially expressed in MN1-Ib and a few additional dorsal Ib MNs. *Mayday* (CG31475) mRNA, encoding a Golgi-localized Ca^2+^ binding protein required for synaptic maintenance of NMJs, was upregulated ∼6-fold in Is. A Mayday Trojan-Gal4 labeled both the dorsal and ventral Is MNs. In contrast, *DIP-η* (>70-fold RNA enrichment in Is) was only expressed in the dorsal Is, suggesting a role in synaptic targeting or maintenance for dorsal muscle NMJs. Together with confirmation of previously known DEGs, cell-type specific labeling indicates Isoform-PatchSeq differentiates Ib and Is transcriptomes. Although mRNA localization and translation rates influence protein production, and post-translational modifications (PTMs) regulate protein localization and turnover, these data provide a foundation for characterizing DEGs that contribute to distinct Is and Ib properties.

### Analysis of Ib and Is MN transcriptomes

To further examine the transcriptional landscapes of Ib and Is, neuronal and synaptic gene classes (Littleton, 2000; Littleton and Ganetzky, 2000; Lloyd et al., 2000) were assembled and analyzed (Supplemental Table 3, Supplemental Item 2). No differences were observed for genes encoding SV proteins and the fusion machinery. Although Is synapses contain larger SVs, endocytosis genes were similarly expressed except for higher Epsin (lqf) and Sorting Nexin 16 (Snx16) expression in Is. For genes encoding AZ proteins, RIM, RBP, and Teneurin-m (Ten-m) were upregulated in Is and previously shown to control AZ organization (Graf et al., 2012; Liu et al., 2011; Mosca et al., 2012). Multiple genes encoding synaptic cleft proteins were differentially expressed, including laminins and several secreted glycoproteins. mRNA for the acetylcholine receptor clustering protein Hig was enriched ∼10-fold in Is, suggesting unique postsynaptic organization. Among ionotropic neurotransmitter receptors, only the *nACHβ2* receptor gene was increased in MN1-Ib. No changes in voltage-gated Ca^2+^ or Na^+^ channel genes or their associated subunits were observed. Several K^+^ channel genes were upregulated in MN1-Ib, including *Shaw, Shab, SK, Ih* and *CNGL*, while *KCNQ* was increased in Is. Genes regulating ion balance and excitability were upregulated in the tonic Ib subclass, including ion exchangers and transporters (*Na^+^ pump β-subunit, Nhe2, Ncc69, Nckx30C, Kcc, vGlut, Eaat1*). More than 20 DEGs encoded G-protein and neuropeptide receptors, indicating MN1-Ib and Is are subject to distinct neuromodulation. ∼35 DEGs encoded proteins implicated in axonal pathfinding and synaptic targeting. Highly enriched pathfinding genes in Is included *DIP-η, DIP-θ, DIP-α, Dpr2, Derailed* (*drl*) and *Fas2*, while MN1-Ib showed enrichment for *DIP-γ, Dpr4, Dpr10, Dpr18, Beat-IIb, Beat-IIIb* and *Kek3*. Components regulating axonal transport (*Arl8, Unc-119* and the kinesins *Unc-104* and *Pav*) and cytoskeletal organization (*Unc-115b, Cdc42, Cib, Mical, Jupiter, Rap1, Shot, α−Tubulin84B, β−Tubulin56D*) were elevated in Is. Given overexpression of Arl8 and Unc-104 increase synaptic cargo delivery in MNs (Goel et al., 2019; Kern et al., 2013; Pack-Chung et al., 2007; Rosa-Ferreira et al., 2018; Vukoja et al., 2018; Zhang et al., 2016), alterations in a rate-limiting axonal transport machinery may enhance cargo delivery across the larger Is axonal arbors.

Although transcription factors (TFs) driving early neuronal differentiation are unlikely to be present at the 3^rd^ instar stage, MN1-Ib and Is expressed unique TFs that may contribute to maintenance of their properties, including *Chinmo, Runt, Spalt major, Luna, Hormone-receptor-like 38*, and *Pangolin* in Ib, and *Trachealess, Huckebein, Zn finger homeodomain 1, Ventral veins lacking, Mamo, Sox14, Cap-n-collar* and *Chip* in Is. Several DEGs encoded signaling proteins in MN1-Ib, including the BMP ligand *Mav*, the wingless ligand *Wnt4*, the FGF ligands *Branchless, Pyramus*, and *Thisbe*, the EFG superfamily ligands *Vein* and *Pvf2*, the Notch ligand *Weary* and the TNF ligand *Eiger*. Downstream components of these signaling pathways were also differentially expressed, with *Toll-6,* the BMP ligand-binding protein *Crimpy*, and the RhoGEF *Still life* (*sif*) upregulated in Ib, and *Dishevelled*, *Myopic*, *Trio*, *RohGEF2* and *Spartin* upregulated in Is. Several mRNAs encoding members of the kinome and phosphatome were differentially expressed, including *CAMKII, PKA-R1, Pkn* and *Cdi* upregulation in Is and enhanced *Pdk1, Lk6, p38a, Cdk4* and *Eip63E* in Ib. These differences indicate Ib and Is engage unique intracellular signaling pathways beyond their distinct synaptic output.

Genes encoding the core transcriptional and translational machinery were similarly expressed. No evidence for distinct energy requirements was identified at the transcriptional level, as mRNAs involved in mitochondrial function and metabolism were similar. Membrane trafficking components were mostly excluded from the DEGs, including SNAREs, Rabs, COPs, TRAPPs, CATCHRs, HOPS, Golgins, ESCRT and Exocyst genes. The Retromer components *Snx3* and *Snx6* were upregulated in Is, along with the ER-Golgi transport regulator *p24-2*, whose mRNA was enriched 20-fold. Several autophagy genes were also increased in Is, including *Atg1* and *Atg4a*. Multiple mRNAs encoding proteins involved in PTMs were differentially expressed, with the *SiaT* sialyltransferase that adds sialic acid to glycoproteins enriched 13-fold enriched in Ib. In contrast, several genes encoding proteins involved in ubiquitination and degradation were upregulated in Is as described below. A matrix of mRNA expression levels for the 190 most robust DEGs in individual Ib and Is neurons is shown in Supplemental Item 3.

### Functional analysis of DEGs in Ib and Is MNs

To begin characterizing the functional relevance of Is and Ib DEGs, synaptic structure was assayed using available mutants and RNAi knockdown with the OK6 pan-MN Gal4 driver. Disruption of multiple DEGs resulted in Ib or Is synaptic growth defects (Figure 4). Is bouton overgrowth was observed in RNAi knockdowns and mutants of several ubiquitin E3 ligases that were enriched in Is MNs (Figure 4A-E, I), including the F-box proteins CG9003 (p<0.0001) and Skp2 (p<0.01), Ube4B (p<0.01), and the E2 conjugating enzyme Ben (p<0.0001). Although these genes are upregulated in Is, they are also expressed in Ib MNs, and disruptions of their function caused a milder overgrowth at Ib NMJs (Figure 4J). Disruptions of the secreted glycoprotein Ana (p<0.01, Figure 2I) and the transport regulator p24-2 (p<0.01, Figure 2I) caused synaptic undergrowth at Is NMJs. Mutations in Kibra (a regulator of Hippo signaling) displayed overgrowth of both Is and Ib NMJs (p< 0.05, Figure 4I, J). Ib-specific synaptic undergrowth was observed in mutants disrupting *SiaT* (p<0.0001, Figure 4G, J), the gene encoding the sole sialytransferase that mediates sialylation of the cell-surface and ECM proteome (Koles et al., 2004). *Wnt4* mRNA was increased 40-fold in Ib MNs, with prior work suggesting a role in synapse specificity (Han et al., 2020; Inaki et al., 2007). *Wnt4* mutants show reduced synaptic growth at Ib terminals (p<0.01, Figure 4H, J), indicating a role in Ib axonal arbor growth.

**Figure 4.**
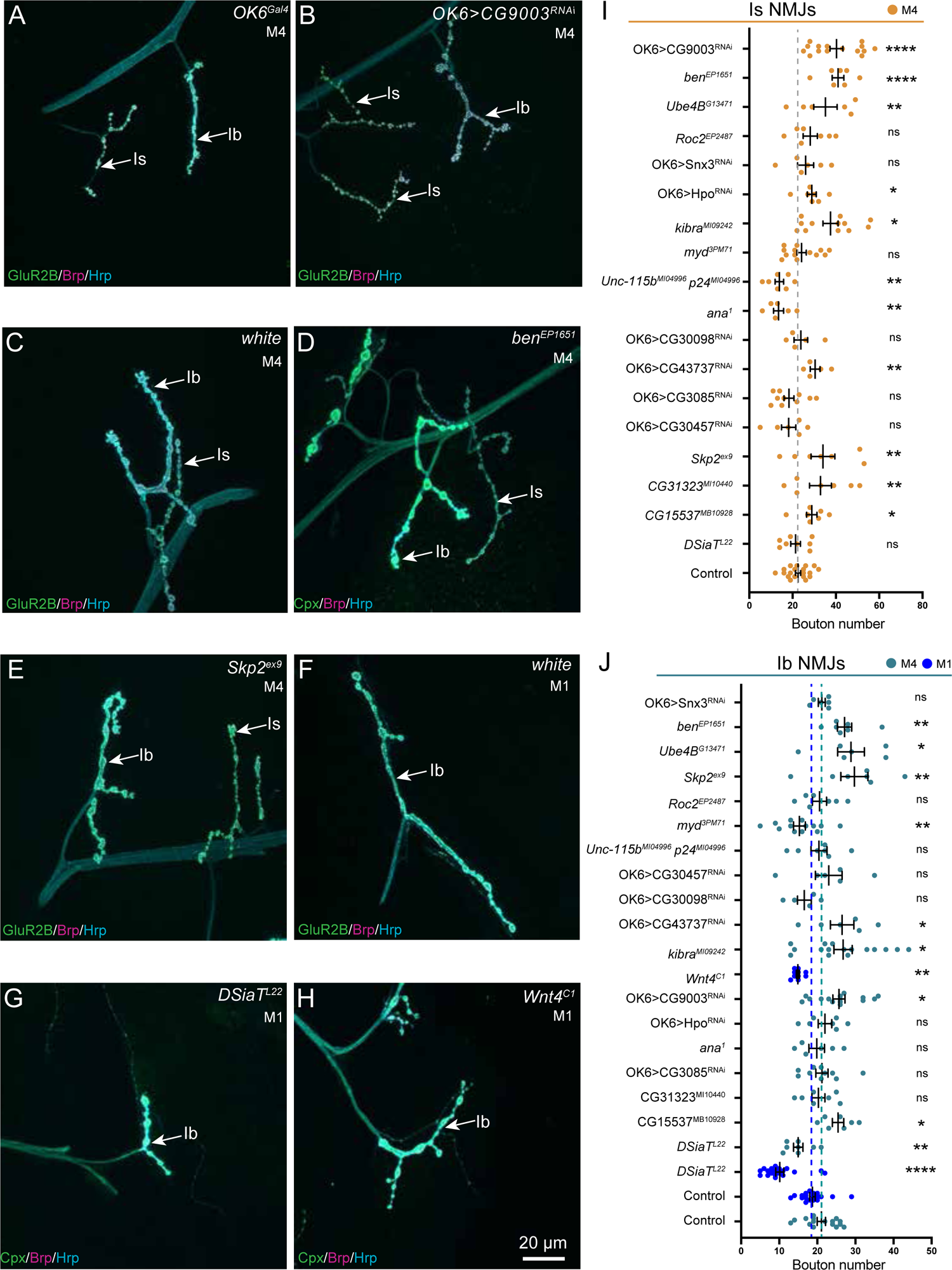
Morphological screen for synaptic growth defects following disruption of DEGs. Representative confocal images of 3^rd^ instar larval NMJs (M1=muscle 1; M4=muscle 4) immunostained for GluR2B or complexin (green), Brp (magenta) and Hrp (cyan) in the indicated genotypes: (A) *OK6 Gal4* driver control line. (B) *OK6>CG9003^RNAi^*. (C) *w*^1118^ control (M4). (D) *ben^EP1651^* mutant. (E) *Skp2^exp^* mutant. (F) *w*^1118^ control (M1). (G) *DSiaT^L22^* mutant. (H) *Wnt4^C1^* mutant. (I) Quantification of Is bouton growth for controls and the indicated mutants or RNAi knockdowns. (J) Quantification for Ib bouton growth for controls and the indicated mutants or RNAi knockdowns. Data represents mean ± SEM (n=6-16 NMJs from 3-8 larvae for all genotypes in I and J). Student t-test, \**p* < 0.05, \*\**p* < 0.01, \*\*\**p* < 0.001, \*\*\*\**p* < 0.0001, ns = not significant.

Given the role of the cytoskeleton in AZ organization (Barber et al., 2018; Blunk et al., 2014; McNeill et al., 2020), STED was used to analyze the nanoscopic structure of Ib and Is AZs following disruption of several DEGs linked to cytoskeletal function. The Is-specific CG3085 protein encodes a *Drosophila* tektin of unknown function, although tektins are enriched in neurons and regulate microtubule stability in other species (Amos, 2008; Norrander et al., 1998). CG3085 RNAi knockdown with OK6-Gal4 disrupted AZ structure in a Is-specific manner (p<0.0001, Supplemental Figure 3A, B). While the donut-like organization of Ib AZs was unchanged, Is AZs were elongated with reduced circularity. The Ib-enriched Toll-6 neurotrophin receptor controls microtubule dynamics at NMJs (McIlroy et al., 2013; McLaughlin et al., 2016). *Toll-6* null mutants displayed altered Ib AZ nanostructure with reduced circularity (p<0.0001, Supplemental Figure 3C, D), suggesting cell-type specific cytoskeletal regulation contributes to tonic and phasic MN development.

### Distinct post-translational modifications regulate diversity in NMJ growth and AZ structure

PatchSeq suggests unique post-translational modifications (PTMs) impact synaptic development in a cell-type-specific manner as several DEGs regulate sialylation and ubiquitination, with disruption to their function altering synaptic growth (Figure 4). Previously described *SiaT* mutants (Repnikova et al., 2010) and CG9003 RNAi knockdown lines (Baccino-Calace et al., 2022; Meltzer et al., 2019) showed reductions in NMJ evoked responses, indicating a functional role for sialylation and ubiquitination. *SiAT* mutants displayed reduced AZ number (p<0.0001, Figure 5A) and synaptic area (p<0.0001, Figure 5B) at Ib NMJs compared to controls or mutant Is NMJs (Figure 5A-C), indicating a preferential role for sialylation in arbor growth of tonic MNs. Beyond CG9003 and Skp2, multiple ubiquitin pathway components were upregulated in Is, including Roc2, Cul5, Posh and the ubiquitin ligases Ube4B, CG2993, CG3014 and LUBEL (Supplemental Item 4), suggesting phasic MNs may employ a specific ubiquitin code to regulate their synaptic properties. Indeed, pan MN knockdown of CG9003, a component of the SCF ubiquitin ligase complex, resulted in increased AZ number (OK6-Gal4 vs OK6>CG9003^RNAi^: p<0.01; CG9003^RNAi^ vs OK6>CG9003^RNAi^: p<0.0001, Figure 5D) and synaptic area (OK6-Gal4 vs OK6>CG9003^RNAi^: p<0.05; CG9003^RNAi^ vs OK6>CG9003^RNAi^: p<0.0001, Figure 5E) in Is MNs, with no effect at Ib NMJs (Figure 5D-F). In mushroom body (MB) neurons, GC9003 functions with the Supernumerary limbs (Slmb) F-box protein to regulate degradation required for dendrite pruning (Meltzer et al., 2019). To determine if CG9003 and Slmb function similarly in regulating Is synaptic growth, *Slmb* RNAi knockdowns were assayed. Like CG9003, *Slmb* knockdown increased Is bouton and AZ number (Supplemental Item 5A-E), indicating the two F-box proteins may regulate axonal growth similar to their role in MB dendrite pruning (Figure 5G).

**Figure 5.**
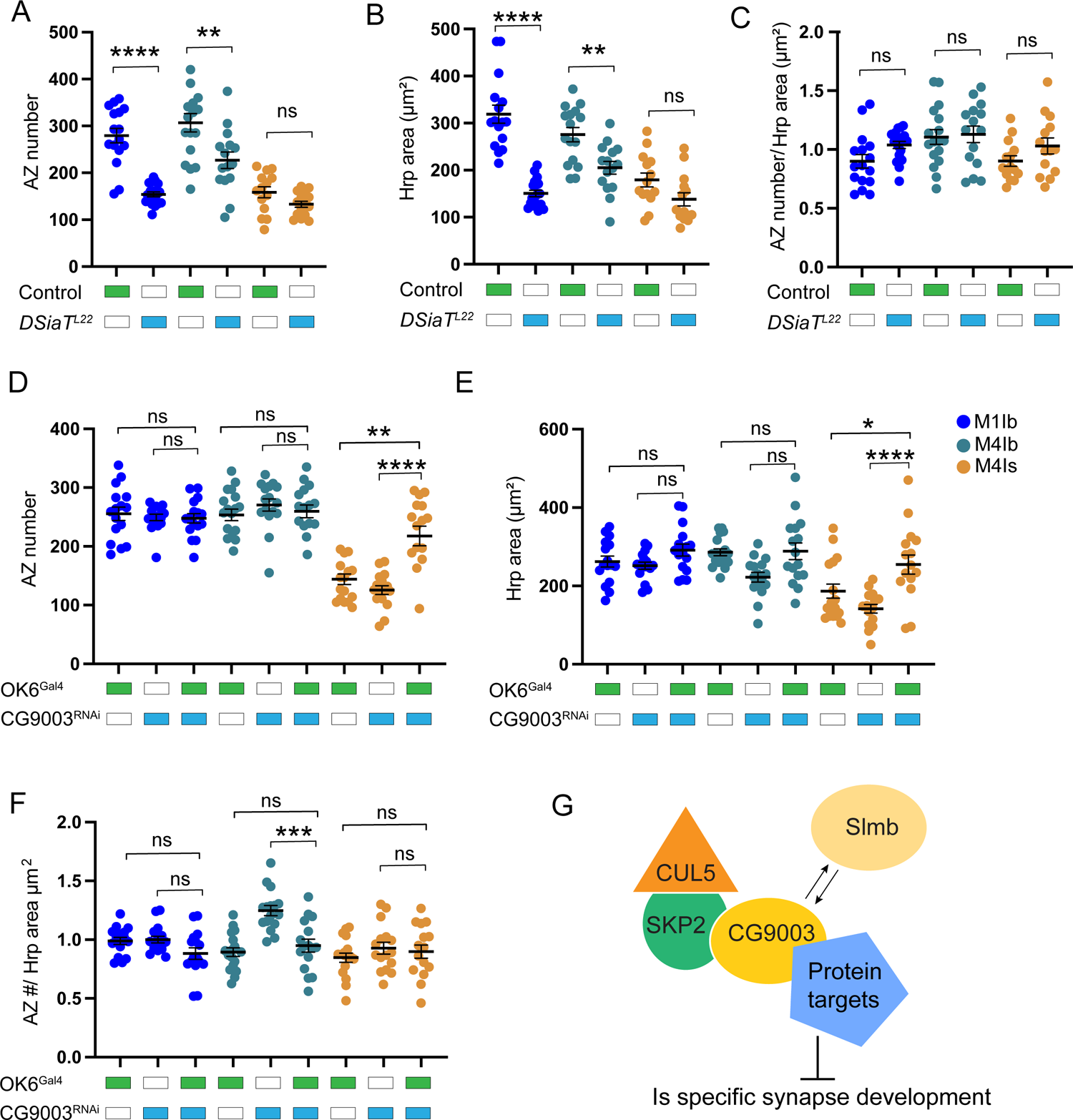
Differential PTMs regulate Ib and Is NMJ growth and AZ number. (A) Quantification of reduced Ib AZ number in *DSiaT^L22^* mutants at muscles 1 and 4 compared to *white* controls. (B) Quantification of reduced Ib synapse area measured by HRP immunostaining in *DSiaT^L22^* at muscles 1 and 4 compared to controls. (C) Quantification of AZ number/area in *DSiaT^L22^* mutants and controls at muscles 1 and 4. (D) Quantification of enhanced Is AZ number in OK6*>*CG9003^RNAi^ compared to controls of OK6-Gal4 driver alone and UAS-CG9003^RNAi^ alone. (E) Quantification of Is synapse area measured by HRP immunostaining in OK6>*CG9003^RNAi^* and controls. (F) Quantification of AZ number/area in OK6>*CG9003^RNAi^* and controls. (G) Model of the SKP2-CG9003-SCF ubiquitin ligase complex and its role in inhibiting Is synaptic growth. Data represents mean ± SEM (n=15 NMJs from 8 larvae for panels A-F). One-way ANOVA with Tukey correction was used to determine significance. \**p* < 0.05, \*\**p* < 0.01, \*\*\**p* < 0.001, \*\*\*\**p* < 0.0001, ns not significant.

To determine if differential PTMs regulate Ib and Is AZ structure, STED imaging was performed in *SiaT* mutants and CG9003 knockdowns. CG9003 knockdown disrupted the triangular organization of Is AZs, causing an increase in circularity (p<0.0001, Figure 6A-C). The nanoscopic organization of Ib AZs was unaltered, indicating CG9003-mediated protein ubiquitination regulates synaptic growth and AZ structure preferentially at Is terminals. In *SiaT* mutants, the donut-like organization of Ib AZs was disrupted with reductions in both AZ area and circularity (p<0.0001, Figure 6D-F), indicating sialylation regulates Ib AZ organization.

**Figure 6.**
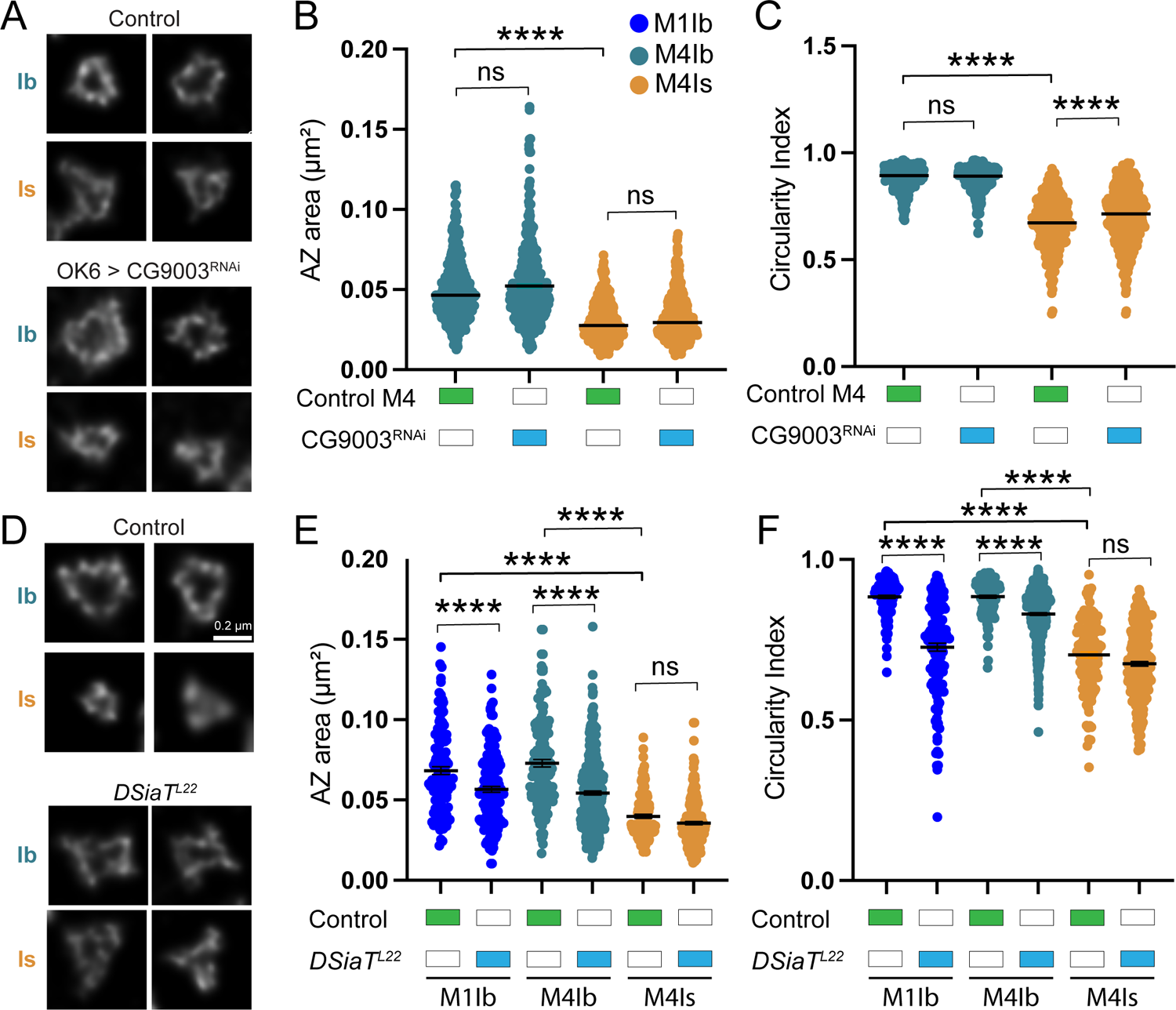
Differential PTMs regulate AZ nanoscopic structure. (A) Representative STED images of Ib and Is AZs in 3^rd^ instar larvae following anti-Brp immunolabeling in OK6-Gal4 controls (upper panels) and OK6-Gal4*>*CG9003 RNAi (lower panels). (B) Quantification of Brp area in control and OK6*>*CG9003^RNAi^. (C) Quantification of circularity index differences between control and OK6>CG9003^RNAi^ (n=363 AZs (control Ib), 428 AZs (OK6>CG9003^RNAi^ Ib), 269 AZs (control Is) and 368 AZs (*OK6>*CG9003^RNAi^ Is), 8 larvae for panels B and C). (D) Representative STED images of Ib and Is AZs following anti-Brp immunolabeling in control (upper panels) and *DSiaT^L22^* mutants (lower panels). (E) Quantification of Brp area differences between control and *DsiaT^L22^*. (F) Quantification of circularity index differences between control and *DSiaT^L22^* (n=124 AZs (control Ib M1), 164 AZs (*DSiaT^L22^* Ib M1), 5 larvae; n= 150 AZs (Control Ib M4), 5 larvae and 340 AZs (*DSiaT^L22^* Ib M4) 8 larvae; n= 132 AZs (control Is M4), 5 larvae, and 234 AZs (*DSiaT^L22^* Is M4), 8 larvae). Data represents mean ± SEM. One-way ANOVA with Tukey correction was used to determine significance. \**p* < 0.05, \*\**p* < 0.01, \*\*\**p* < 0.001, \*\*\*\**p* < 0.0001, ns not significant.

### Calbindin 53E contributes to resting [Ca^2+^] differences at Ib and Is synapses

Cbp53E encodes a six EF-hand containing protein and is the *Drosophila* homolog of vertebrate calbindin and calretinin proteins that function as Ca^2+^ buffers in the nervous system. Prior studies indicated Cbp53E regulates MN axonal branching (Hagel et al., 2015) and its downregulation contributes to Ca^2+^ defects in *Drosophila* Fragile X models (Tessier and Broadie, 2011). Given Cbp53E is enriched 30-fold in Is versus Ib MNs, and Is AZs have ∼50% more presynaptic Ca^2+^ influx following nerve stimulation (Figure 1J), differences in Cbp53E-mediated Ca^2+^ buffering might contribute to Is specific Ca^2+^ regulation. To test this hypothesis, Ca^2+^imaging was performed in control and *Cbp53E* mutants using AZ-targeted Brp-GCaMP7s (Figure 7A). Raw Brp-GCaMP7s fluorescence (Figure 7B) and Brp-GCaMP7s AZ fluorescence normalized to AZ Brp levels determined post-fixation following Brp immunostaining (Figure 7C) were acquired. Resting Ca^2+^ levels were 27% higher at Ib AZs compared to Is in controls (p<0.0001). The difference in resting [Ca^2+^] between the two MN populations was abolished in *Cbp53E* mutants, which showed no change in [Ca^2+^] at Ib AZs and enhanced resting [Ca^2+^] in Is (Con Is vs *Cbp53E* Is: p<0.001; *Cbp53E* Ib vs *Cbp53E* Is: p=0.94, Figure 7B, C)). These data indicate differences in Cbp53E expression between tonic and phasic MN populations plays a role in setting baseline [Ca^2+^] levels.

**Figure 7.**
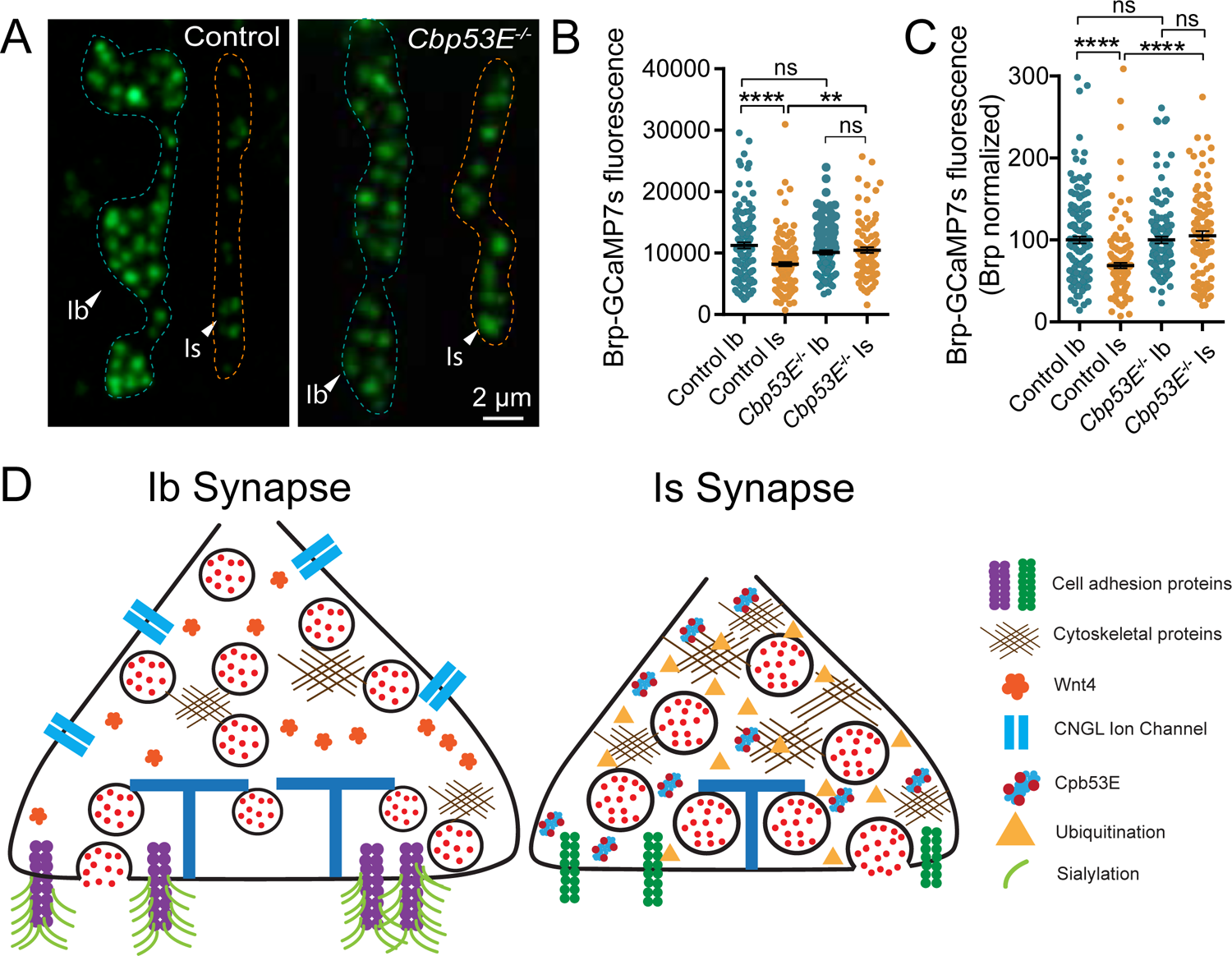
Increased resting Ca^2+^ in *Cbp53E* mutants and synaptic models for Ib and Is MNs. (A) Representative images of resting Ca^2+^ fluorescence at Ib and Is NMJ AZs at muscle 4 in control and *Cbp53E* mutant larvae expressing Brp-GCaMP7s. (B) Quantification of resting Ca^2+^ levels at Ib and Is AZs at muscle 4 in control and *Cbp53E* mutants expressing Brp-GCaMP7s. (C) Quantification of resting Ca^2+^ levels normalized to AZ Brp abundance at Ib and Is AZs at muscle 4 in control and *Cbp53E* mutants expressing Brp-GCaMP7s (n=135 AZs (control Ib), 139 AZs (control Is) from 8 NMJs, 3 larvae; Is: n=127 AZs (*Cbp53E* Ib) from 7 NMJs, 95 AZs (*Cbp53E* Is) from 6 NMJs, 3 larvae for panels B and C). One-way ANOVA with Tukey correction was used to determine significance. \**p* < 0.05, \*\**p* < 0.01, \*\*\**p* < 0.001, \*\*\*\**p* < 0.0001, ns not significant. (D) Model of Ib and Is synapses highlighting key differences identified in this study.

## Discussion

In the current study we describe nanoscopic, transcriptomic, and post-translational signatures that contribute to distinct features of tonic and phasic MN synapse diversity. While RNAseq has generated expression profiles for many neuronal subtypes, the relevant DEGs that drive functional and structural diversity are poorly understood. We used single neuron Isoform-PatchSeq to determine transcriptional profiles of two well-studied glutamatergic MN subtypes in *Drosophila* and carried out genetic analysis to examine the role of several DEGs in specifying synaptic properties. We identified a distinct compact AZ structure and enhanced Ca^2+^ entry at phasic Is synapses. Our transcriptomic analysis identified ∼800 DEGs that may support key differences in neuronal properties between these two MN subtypes, including Cbp53E that regulates resting Ca^2+^ levels, the cytoskeletal regulators CG3085 and Toll-6 that contribute to AZ nanostructure, and the Wnt4 ligand that participates in MN-specific synaptic growth programs. In addition, DEGs controlling sialylation and ubiquitination altered synaptic growth and AZ organization in a MN-specific manner. Genes encoding neuropeptide receptors, synaptic cleft proteins, axonal pathfinding and cell adhesion molecules, membrane excitability and ionic balance regulators, and the Arl8 and Unc-104 axonal transport factors, were also differentially expressed. Together, these data provide a framework for characterizing the molecular mechanisms contributing to unique features of tonic and phasic synapse diversity (Figure 7D).

Our analysis is consistent with a model that AZ structural differences and altered Ca^2+^ entry and buffering contribute to release differences between Ib and Is MNs. Given phasic Is synapses have enhanced synaptic strength and higher *Pr*, their unique stellate AZ architecture and smaller cytomatrix, together with shorter T-bars and larger SVs, could contribute to distinct AZ Ca^2+^ dynamics and alterations in coupling Ca^2+^ influx to SV fusion. At tonic synapses, reduced Ca^2+^ influx and SV fusogenicity would favor lower *Pr*. Previous studies demonstrated changes in the coupling distance between SVs and Ca^2+^ channels driven by distinct AZ components can alter synaptic strength at a number of synapse types (Bucurenciu et al., 2008; Ghelani and Sigrist, 2018; Nakamura et al., 2015). Increased expression of the Cbp53E Ca^2+^ buffer in Is MNs lowers resting [Ca^2+^], which together with enhanced Ca^2+^ influx, provides a synergistic mechanism with the more compact Is AZ nanostructure that could enhance release. Prior studies identified additional differences in SV availability and fusogenicity in Ib and Is MNs also contribute to differences in their release properties. Reduced synaptic expression of Brp, the decoy SNARE Tomosyn, and the fusion clamp Complexin at Is terminals alters the size of SV pools and their availability for fusion (Jorquera et al., 2012; Mrestani et al., 2021; Newman et al., 2022; Sauvola et al., 2021). These data indicate Ib synapses have fewer fusogenic SVs available to respond to Ca^2+^ entry that also contributes to differences in synaptic output at tonic terminals. No changes in Brp, Tomosyn or Complexin mRNA levels were detected in Isoform-PatchSeq, suggesting post-translational degradation or altered trafficking is likely to underlie differences in their abundance at Is synapses.

Multiple studies indicate PTMs such as ubiquitylation, glycosylation, phosphorylation, acetylation, and sumoylation impact synapse organization and function by modifying protein stability and localization (Cho et al., 2015; Jeoung et al., 2022; Kawabe and Brose, 2011; Mabb, 2021; Miśkiewicz et al., 2011; Scott and Panin, 2014). How distinct PTMs control synapse function in a cell-type-specific manner is poorly understood. Isoform-PatchSeq indicates sialylation and ubiquitination are differentially regulated in Ib and Is MNs, contributing to their unique synaptic properties. Disruptions of other E3 ubiquitin ligases like Highwire alter synaptic growth at *Drosophila* NMJs, though they impact both Is and Ib terminals (Collins et al., 2006; DiAntonio and Hicke, 2004; DiAntonio et al., 2001). Identifying targets of the Skp2-CG9003 E3 ligase complex in Is MNs should clarify mechanisms by which cell-type specific proteolysis contributes to their unique synaptic morphology and function. Several studies also support a key role for sialylation in organization of the cell-surface proteome, regulation of cell-adhesion, and control of neuronal excitability (Ednie and Bennett, 2012; Scott and Panin, 2014). Lack of sialylation impairs surface expression of multiple ion channels and cell adhesion proteins, indicating defects in Ib synaptic growth and AZ organization in *SiaT* mutants could be secondary to a lack of sialylation of multiple cell-surface proteins. Further studies will be required to identify the specific sialylated targets controlling Ib-specific synaptic organization. In summary, these findings highlight multiple pathways and PTMs that contribute to synaptic diversity between tonic and phasic MNs, while providing a resource to characterize additional DEGs that specify other structural and functional features of these unique neuronal subtypes. Neurons with tonic and phasic output represent an evolutionarily conserved design principle across species, and it will be interesting to determine if similar DEGs are involved in specifying their properties. Recent RNA profiling indicates *Drosophila* visual neurons differentially express sialyltransferases and proteolytic regulators (Kurmangaliyev et al., 2020), while mammalian cortical GABAergic interneuron subtypes differentially express sialyltransferases that alter their cell surface proteome (Paul et al., 2017). As such, cell type-specific PTMs may represent a broadly used mechanism to generate neuronal diversity beyond their role in *Drosophila* tonic and phasic MNs.

## Acknowledgements

This work was supported by NIH grants MH104536 and NS117588 to J.T.L. and F31NS118948 to A.B.C. K.L.C. and N.A.S were supported in part by NIH pre-doctoral training grant T32GM007287. We thank the Bloomington *Drosophila* Stock Center (NIH P40OD018537), the Developmental Studies Hybridoma Bank, the Vienna *Drosophila* Resource Center (Austria), Vladislav Panin (Texas A&M), Charles Tessier (Indiana School of Medicine), and Daniel Babcock (Lehigh University) for providing *Drosophila* strains and antibodies, Dina Volfson (MIT) for cloning assistance, Stuart Levine (MIT) for RNAseq library and sequencing assistance, Harvard Imaging Core for STED access, and members of the Littleton lab for helpful discussions and comments on the manuscript.

## Abbreviations

NMJ: neuromuscular junction

SEM: standard error of the mean

WT: wildtype

HRP: horse radish peroxidase

AZ: active zone

MN: motoneuron

*Pr*: release probability

EM: electron microscopy

STED: stimulated emission depletion nanoscopy

DEGs: differentially expressed genes

SV: synaptic vesicle

VNC: ventral nerve cord

GluR: glutamate receptor

VGCC: voltage-gated Ca^2+^channel

TF: transcription factor.

## Supplemental Figure Legends

**Supplemental Figure 1.**
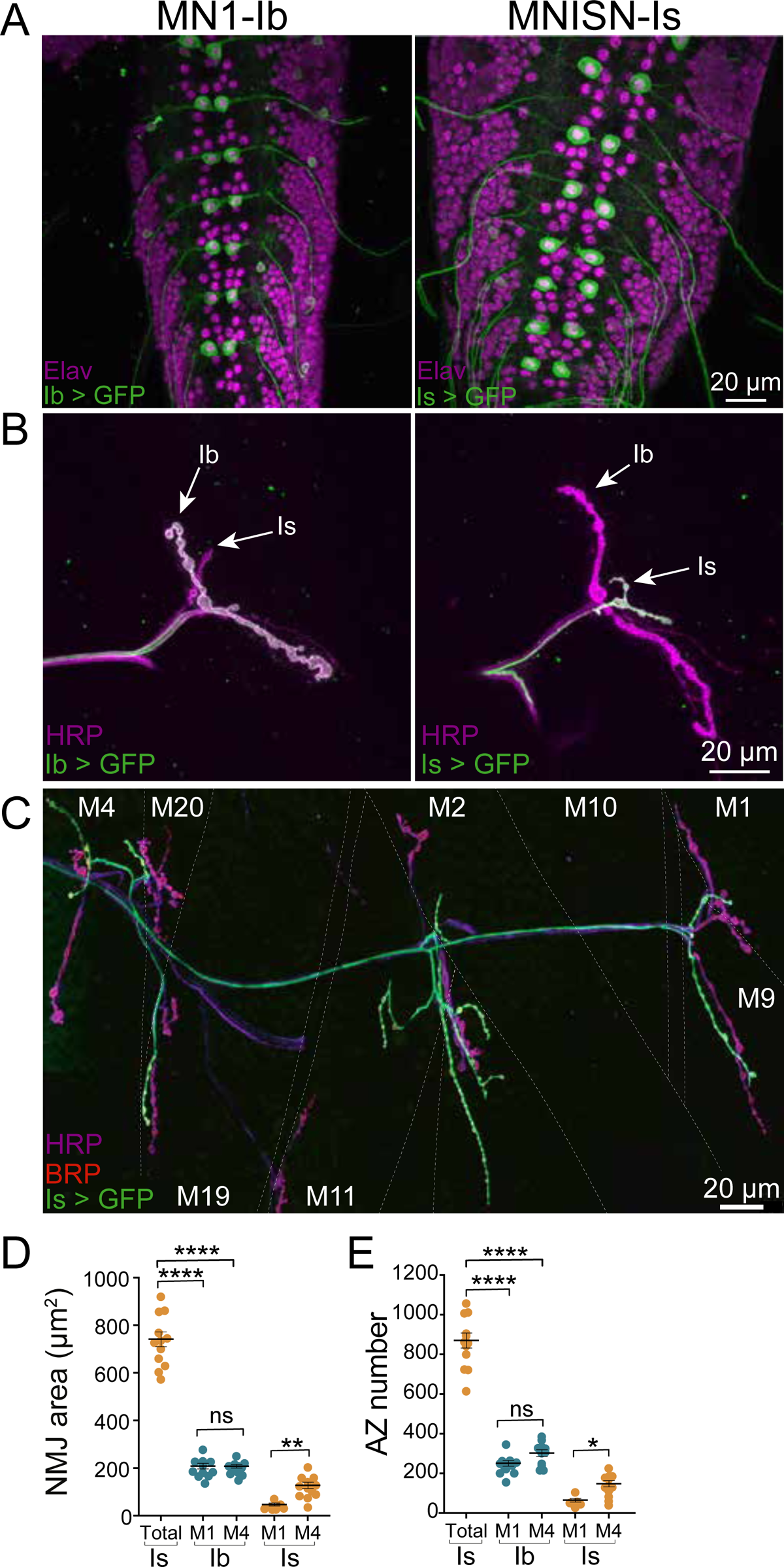
Ib and Is-specific Gal4 driver expression and synaptic quantification. (A) Representative confocal images of 3^rd^ instar larval VNCs of MN1-Ib or Is Gal4 driver lines expressing UAS-GFP and immunostained for the pan-neuronal nuclear protein elav. (B) Representative confocal images of 3^rd^ instar larval NMJs of MN1-Ib or Is Gal4 driver lines expressing UAS-GFP and immunostained for HRP. Ib and Is NMJs are denoted with white arrows. (C) Representative confocal image of a 3^rd^ instar abdominal hemi-segment showing MNISN-Is innervation of the dorsal muscle field in Is Gal4 driver larvae expressing UAS-GFP and immunostained with Brp and Hrp. (D) Quantification of Ib and Is NMJ area determined by HRP staining in 3^rd^ instar larval abdominal segments 3 and 4 for the indicated muscle targets (MN1-Ib versus Is at muscle 1: p<0.0001; MN1-Ib versus Is over all innervated Is muscles: p<0.0001, n=11 NMJs from 6 larvae). (E) Quantification of Ib and Is AZ number determined by Brp staining in 3^rd^ instar larval segments 3 and 4 for the indicated muscle targets (MN1-Ib versus Is at muscle 1: p<0.0001; MN1-Ib versus Is over all innervated Is muscles: p<0.0001, n=11 NMJs from 6 larvae). Four muscle 1 NMJs lacked Is innervation in the data set and were excluded from Ib versus Is AZ number comparison.

**Supplemental Figure 2.**
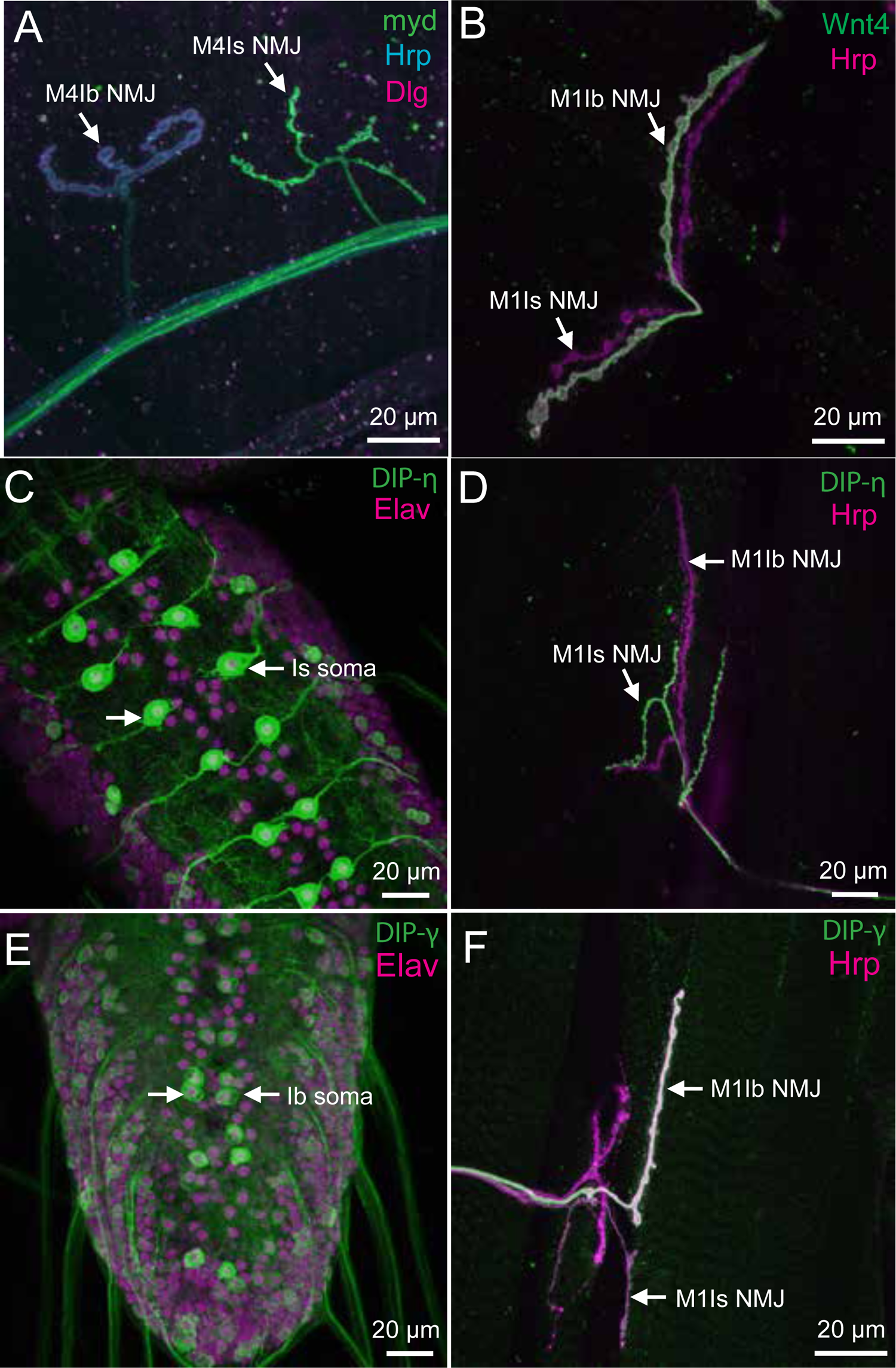
Verification of protein expression differences for several DEGs. (A) Representative confocal image of a 3^rd^ instar larval muscle 4 NMJ of a Trojan Mayday-Gal4 strain driving GFP and immunostained for Dlg and Hrp. White arrows denote Ib and Is NMJs. (B) Representative confocal image of a 3^rd^ instar larval muscle 1 NMJ of a Trojan Wnt4-Gal4 line driving GFP and immunostained for Hrp. White arrows denote Ib and Is NMJs. (C) Representative confocal image of a 3^rd^ instar VNC of a Trojan DIP-η-Gal4 strain driving GFP and immunostained for Elav. White arrows denote the MNISN-Is soma within the VNC. (D) Representative confocal image of a 3^rd^ instar larval muscle 1 NMJ of a Trojan DIP-η-Gal4 strain driving GFP and immunostained for Hrp. White arrows denote Ib and Is NMJs. (E) Representative confocal image of a 3^rd^ instar VNC of a Trojan DIP-γ-Gal4 strain driving GFP and immunostained for Elav. White arrows denote the MN1-Ib soma within the VNC. (F) Representative confocal image of a 3^rd^ instar larval muscle 1 NMJ of a Trojan DIP-γ-Gal4 strain driving GFP and immunostained for Hrp. White arrows denote Ib and Is NMJs.

**Supplemental Figure 3.**
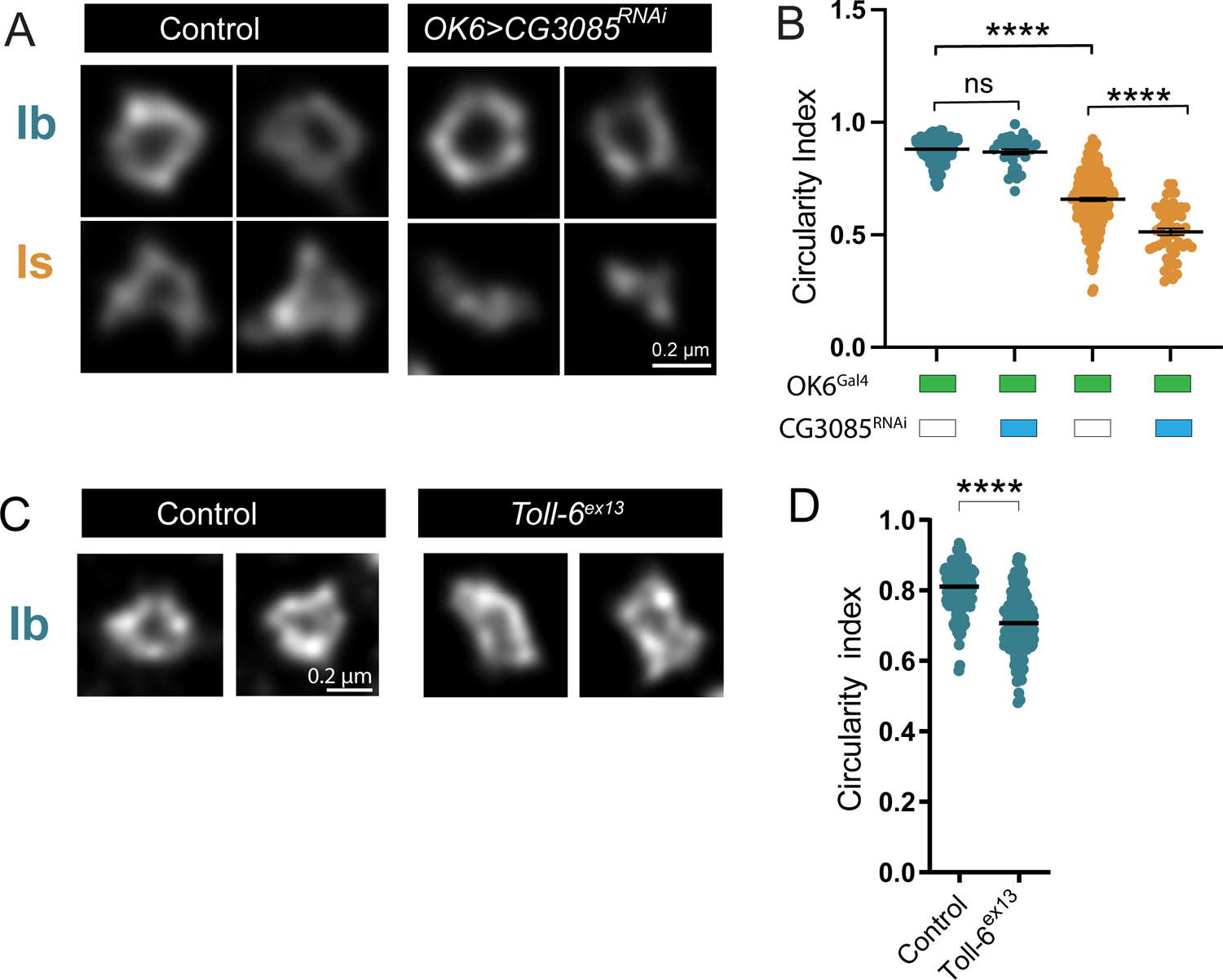
Changes in AZ nanoscopic structure following disruptions of CG3085 and Toll-6. (A) Representative STED images of Ib and Is AZs in 3^rd^ instar larvae following anti-Brp immunolabeling in OK6-Gal4 controls (left panels) and OK6-Gal4*>*CG3085^RNAi^ (right panels). (B) Quantification of circularity index in control and OK6>CG3085^RNAi^ AZs (n=273 AZs (control Ib), 32 AZs (OK6>CG3085^RNAi^ Ib); n= 229 AZs (control Is) and 53 AZs (OK6>CG3085^RNAi^ Is), p<0.0001). (C) Representative STED images of Ib and Is AZs in 3^rd^ instar larvae following anti-Brp immunolabeling in white controls (left panels) and *Toll-6^ex13^* mutants (right panels). (D) Quantification of circularity index in control (n=172 Ib AZs) and *Toll-6^ex13^* (n=192 Ib AZs, p<0.0001). One-way ANOVA with Tukey correction was used to determine significance. \**p* < 0.05, \*\**p* < 0.01, \*\*\**p* < 0.001, \*\*\*\**p* < 0.0001, ns not significant.

---

**Supplemental Table 1.** Full TPM and read counts for all genes in the Isoform PatchSeq dataset from 3^rd^ instar larval MN1-Ib, MNISN-Is, muscle 1 and muscle 4 samples.

**Supplemental Table 2.** Splice isoforms detected for all genes in the Isoform PatchSeq dataset from 3^rd^ instar larval MN1-Ib, MNISN-Is, muscle 1 and muscle 4 samples.

**Supplemental Table 3.** Gene class analysis of Isoform PatchSeq dataset for 3^rd^ instar larval MN1-Ib, MNISN-Is, muscle 1 and muscle 4 samples.

## Supplemental Item Legends

**Supplemental Item 1.**
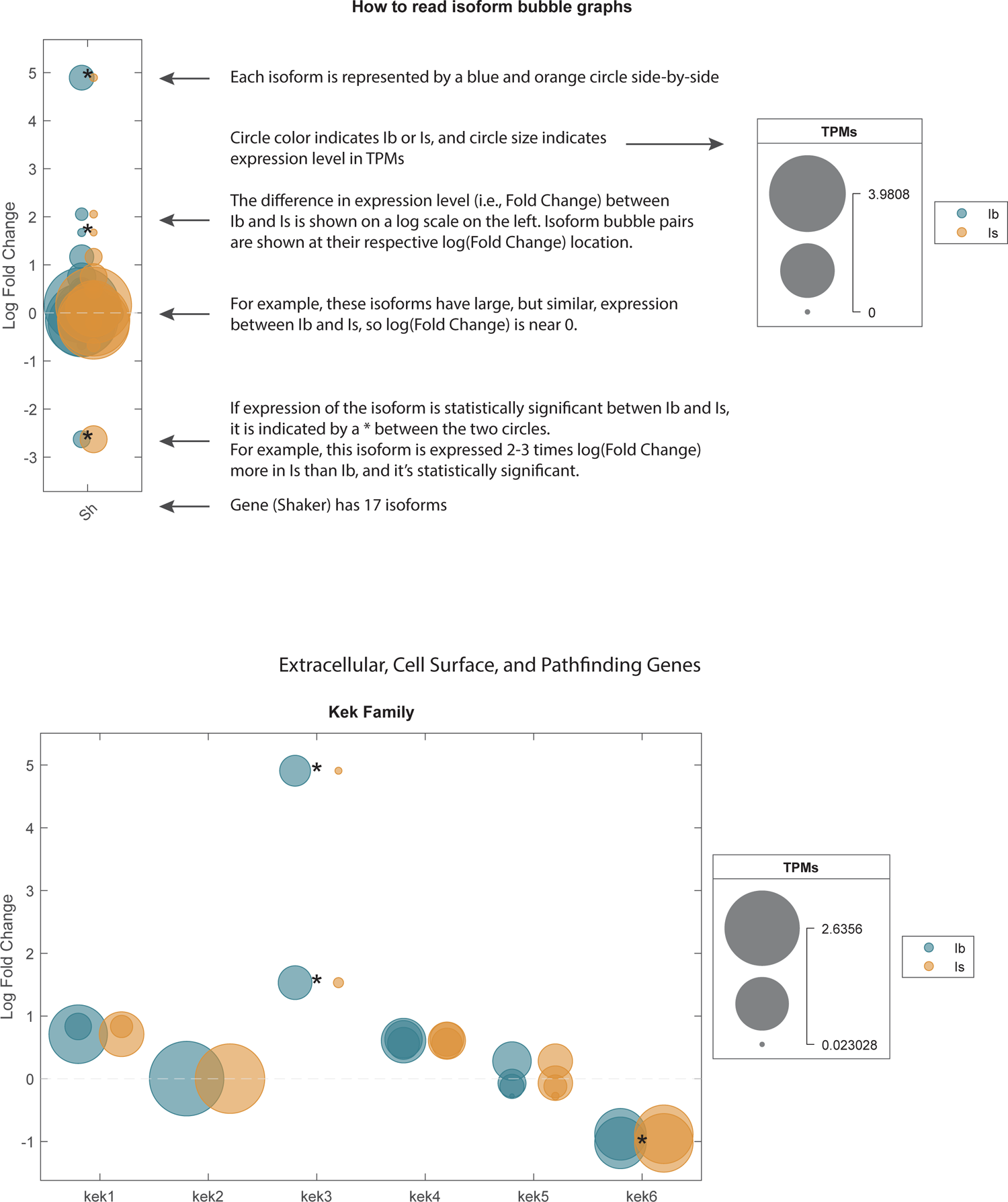
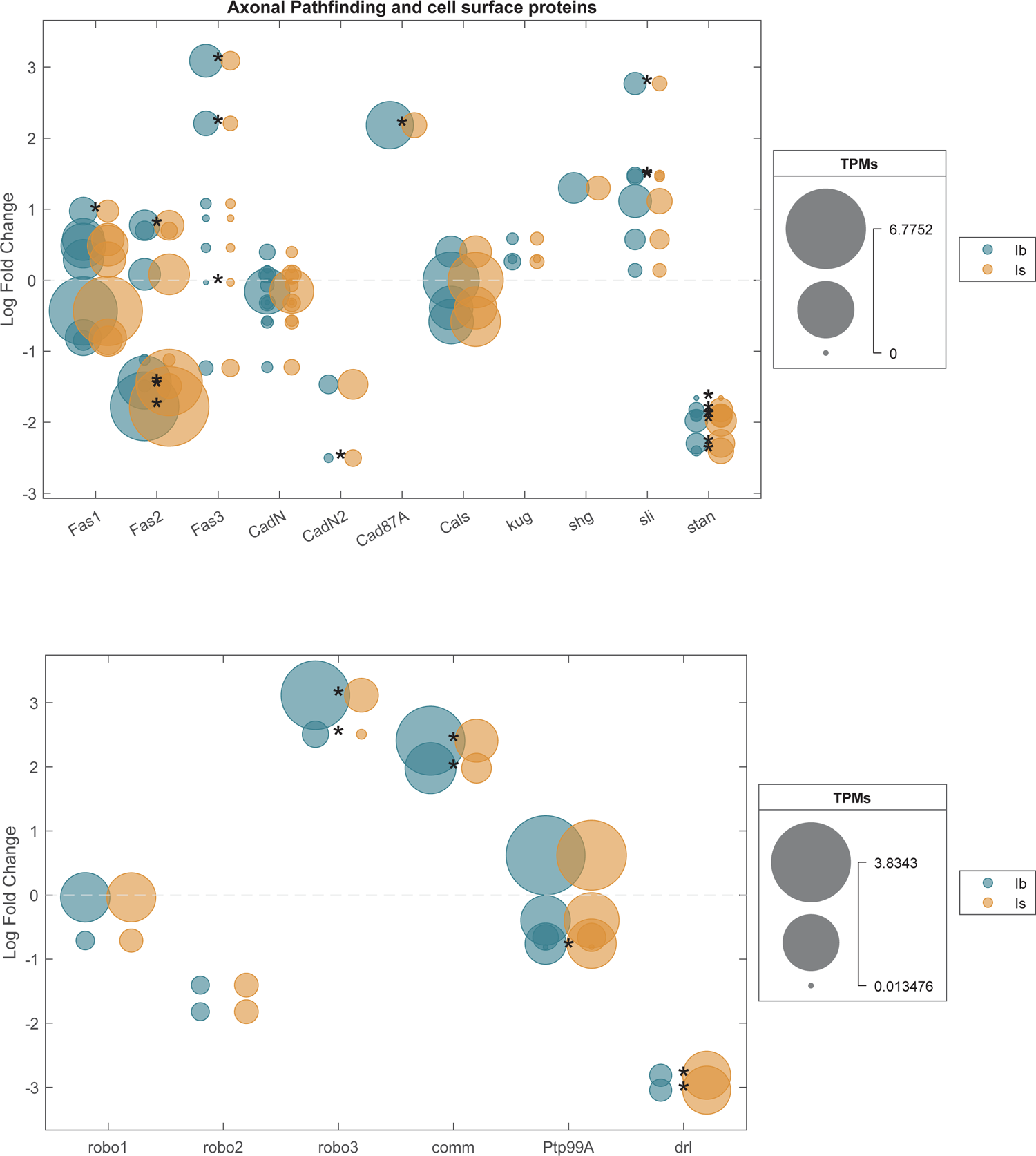
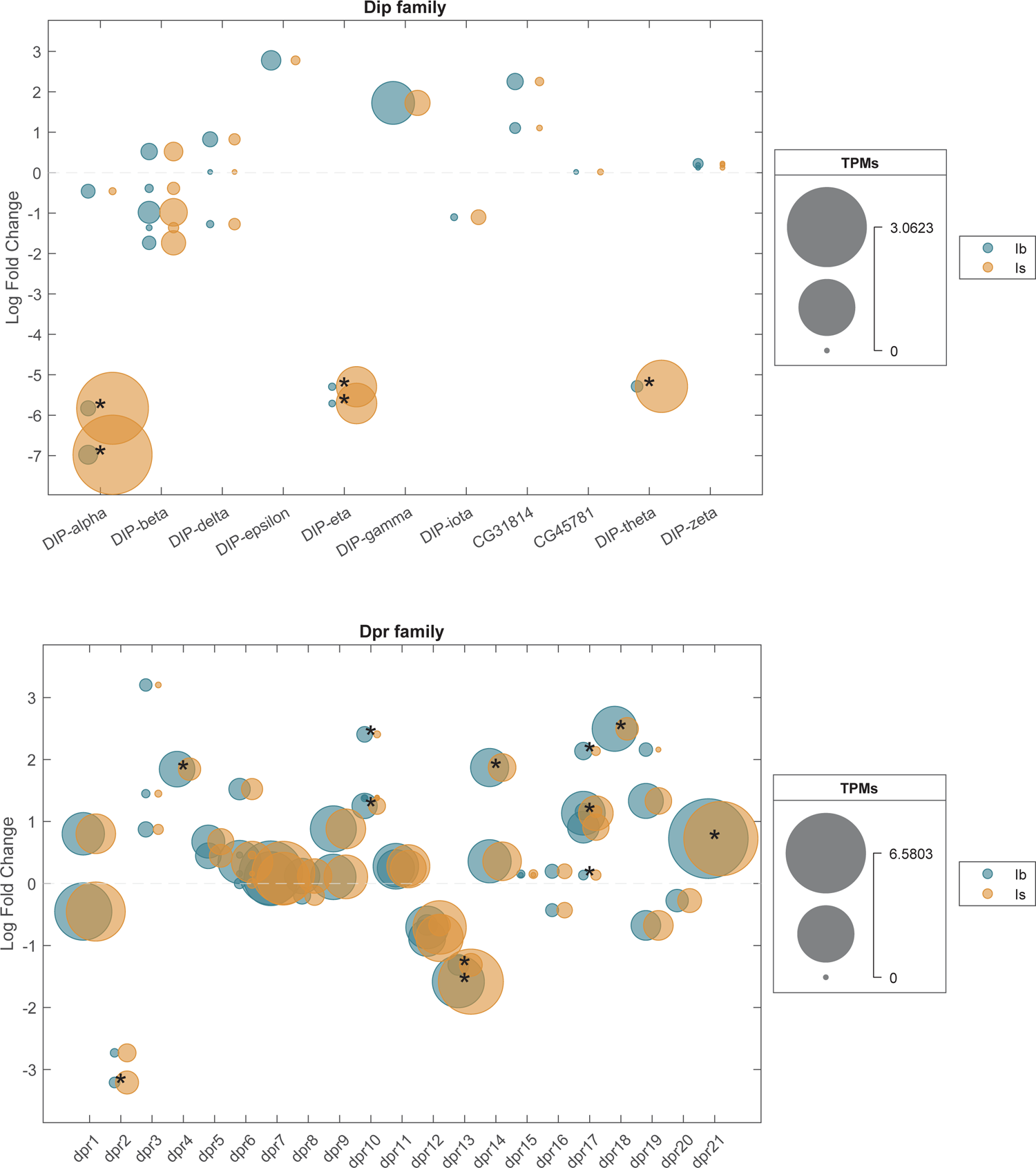
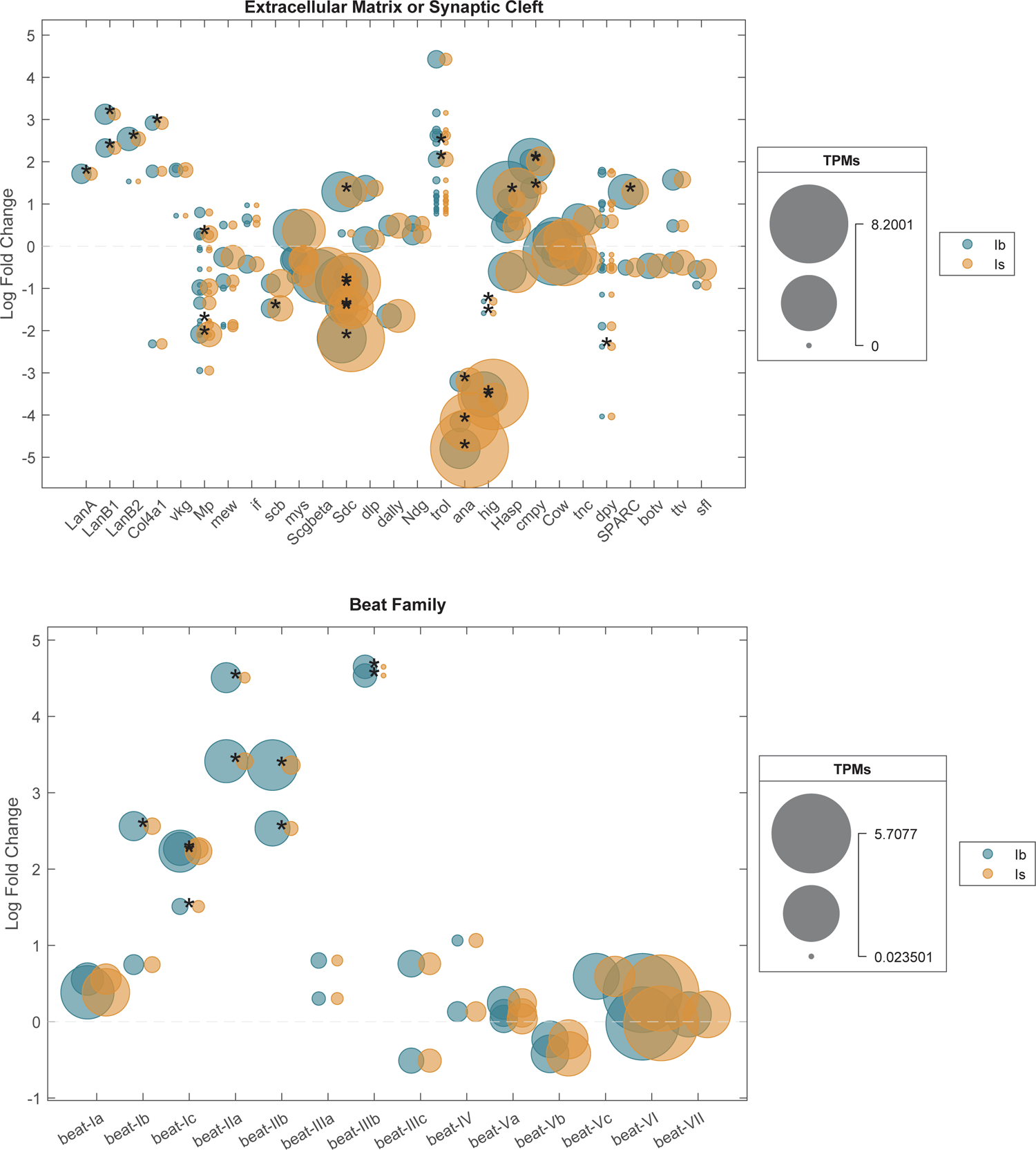
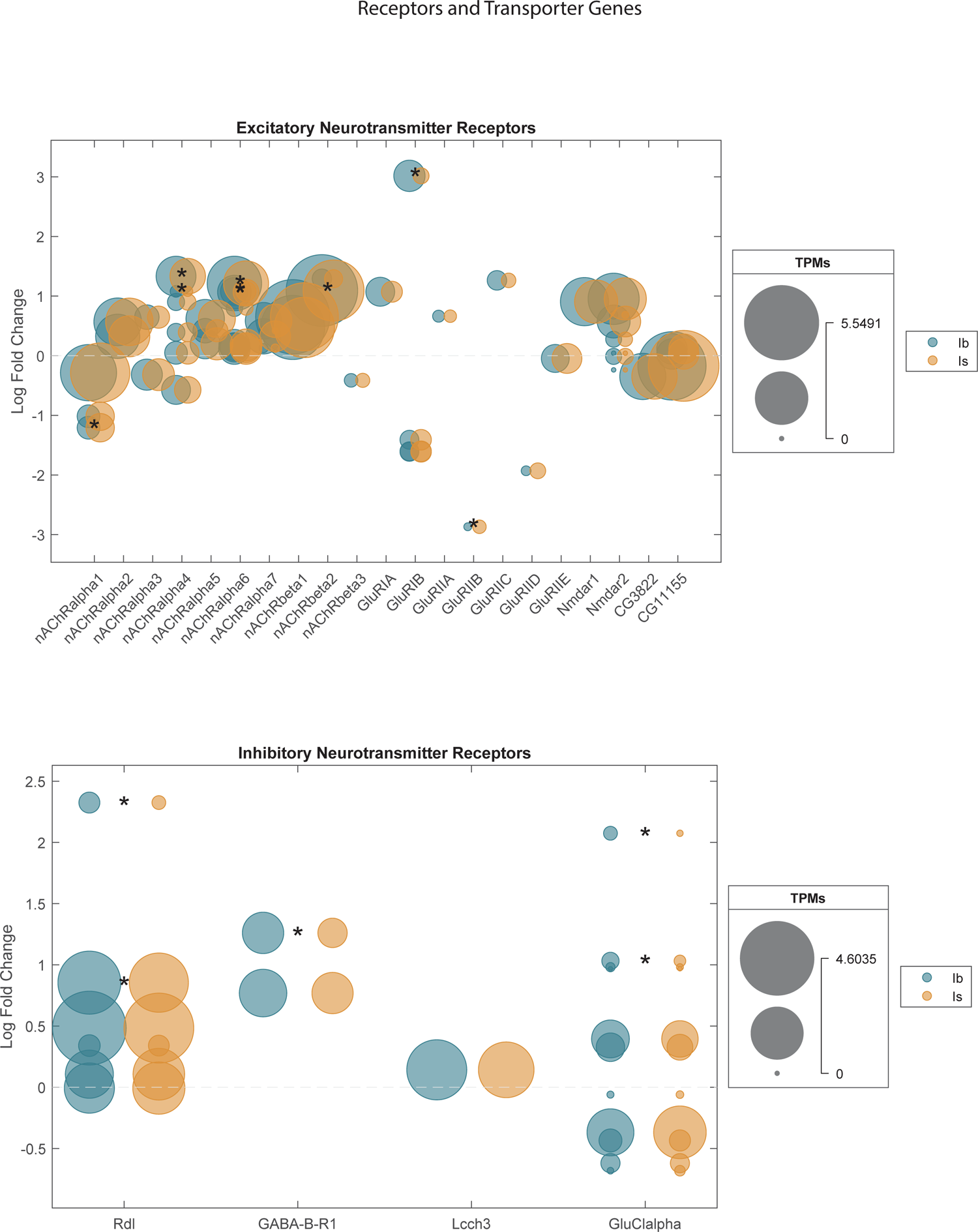
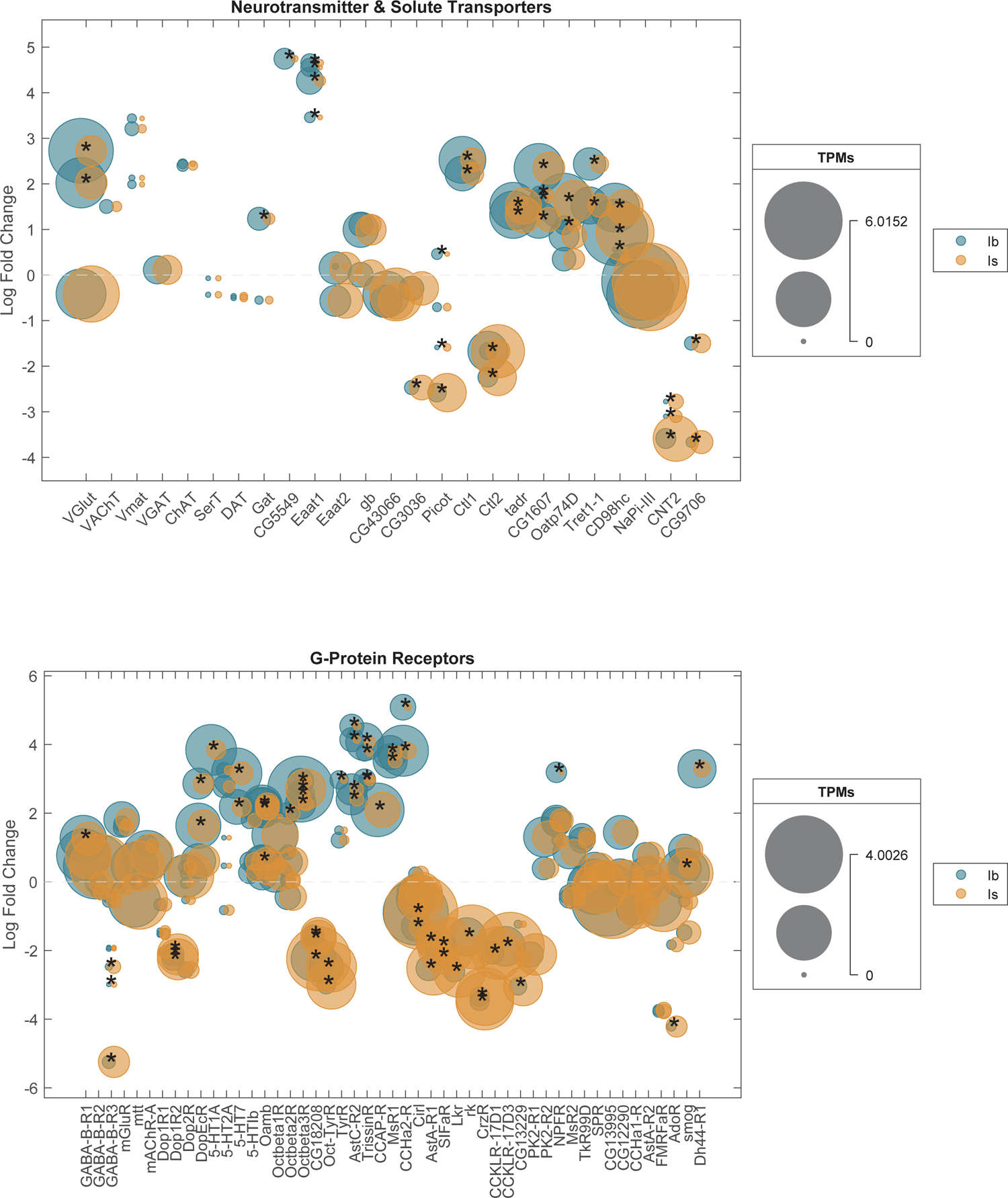
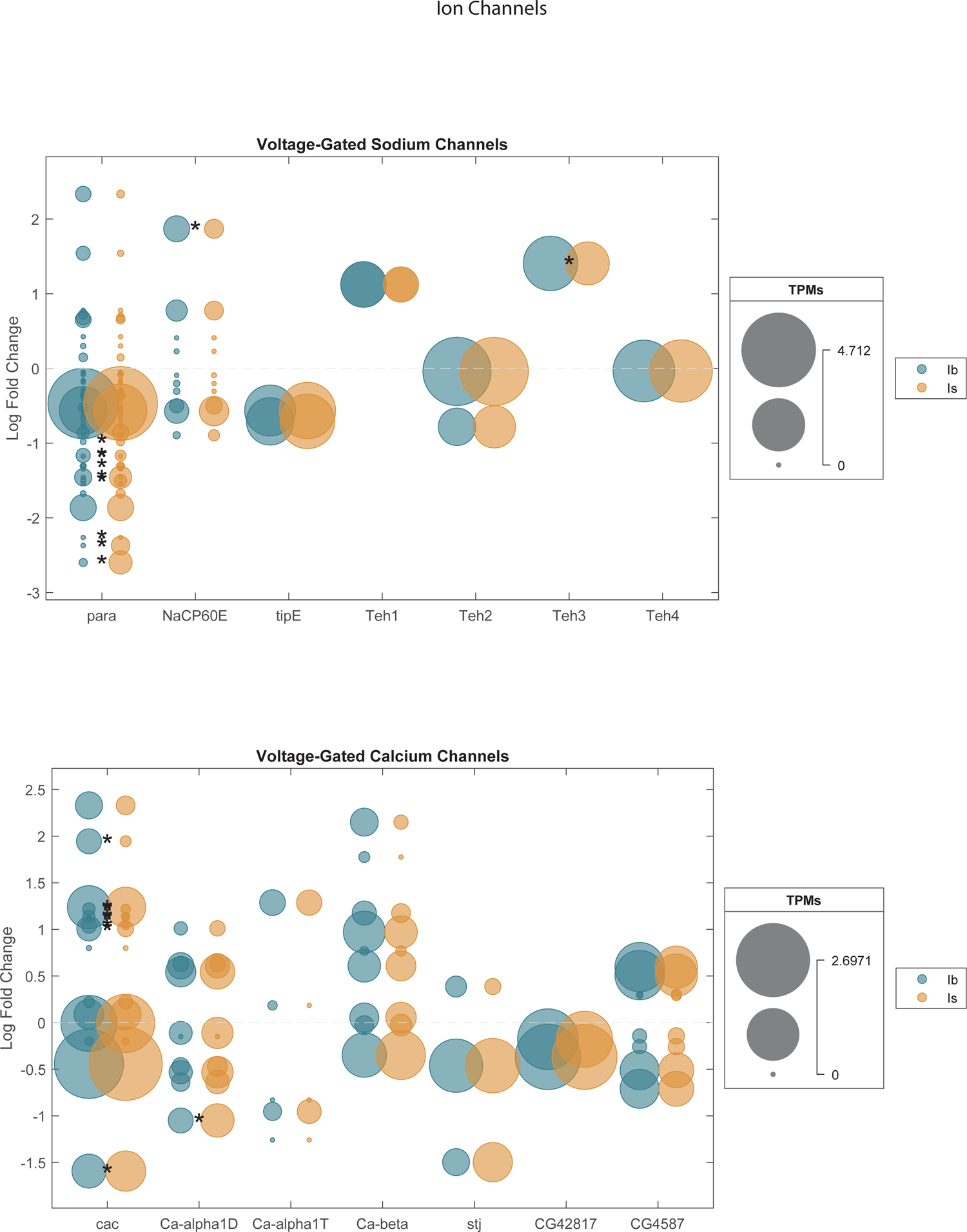
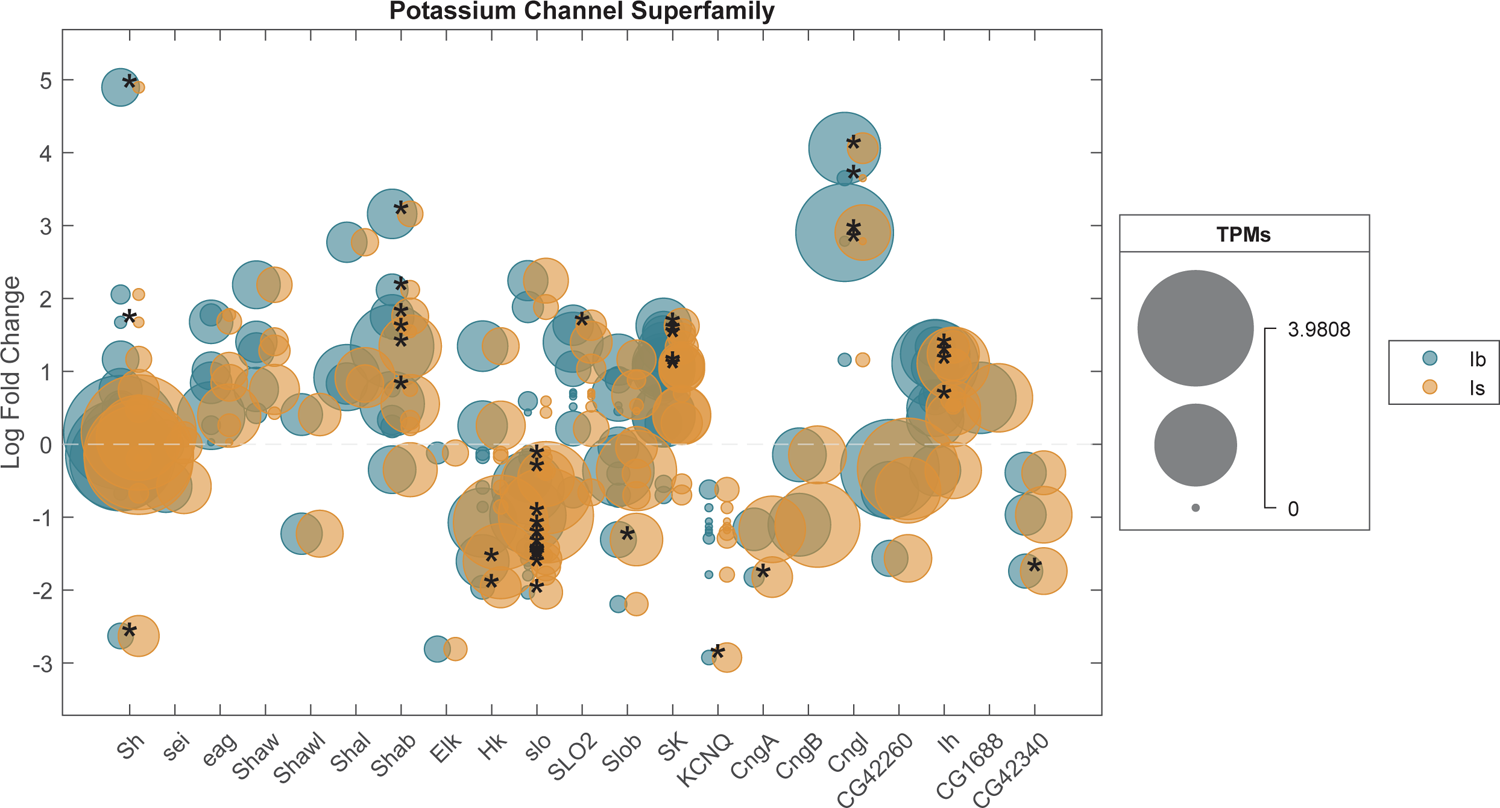
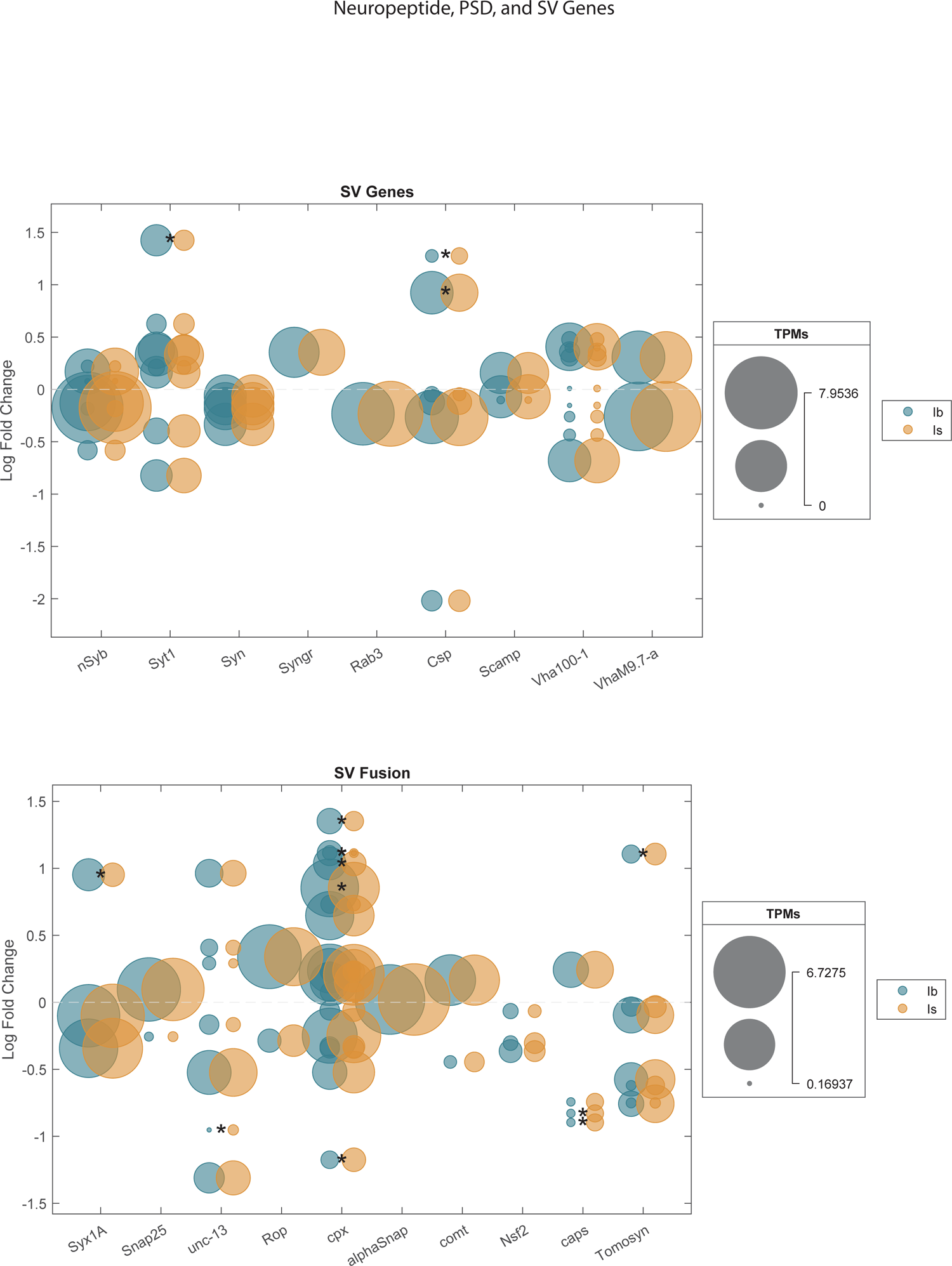
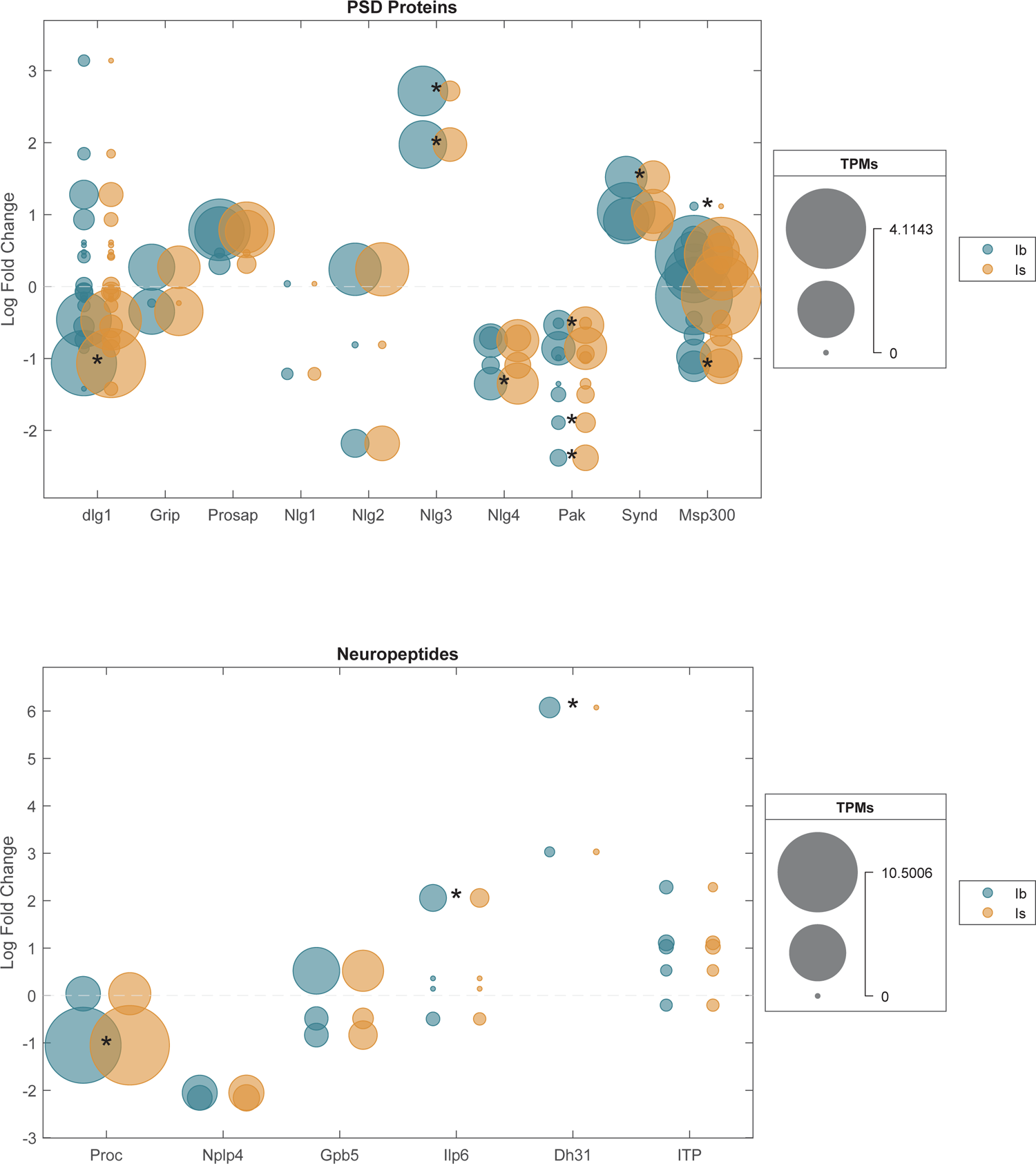
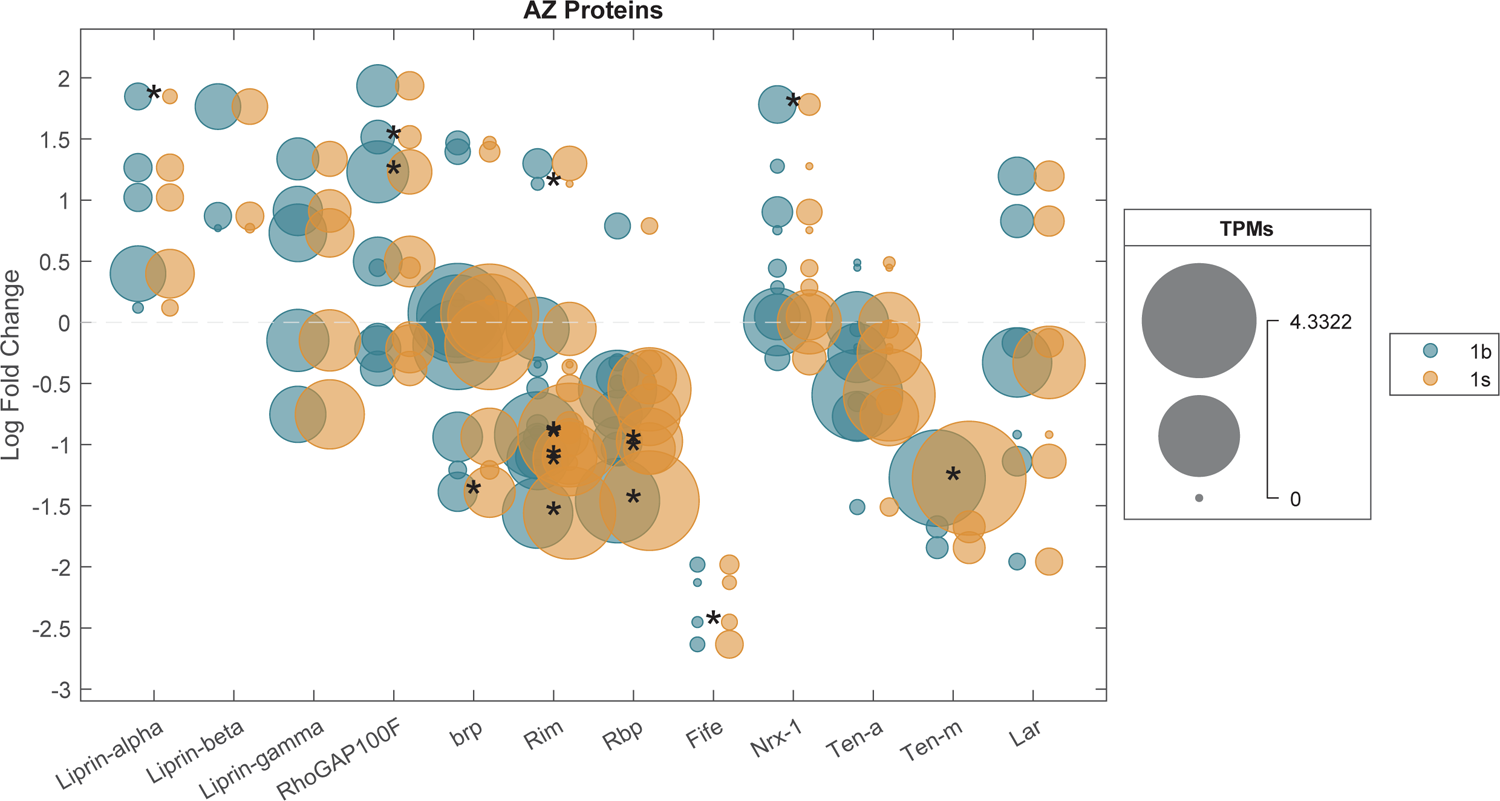
Highlights of splice isoform differences across subsets of neuronal genes. Genes were manually grouped into functional categories. For each gene in a specific category, a pair of blue and orange circles are arranged vertically above the gene’s name. Each pair of circles represents an individual spice isoform from the indicated locus. Circles size indicates expression levels in TPMs (transcripts per million). Small circles indicate lower expression and larger circles represent higher expression. The vertical position of the pair indicates whether that isoform is expressed at higher levels in Ib (positive log fold change) or Is MNs (negative log fold change). The average expression level in TPMs of each isoform in Ib (blue) or Is (orange) cells is indicated by the size of each circle. If the difference in expression between Ib and Is is significant (adjusted p-value from the RNA sequencing Wald statistic), it is indicated by a single asterisk between the orange and blue circles. Numerical data for every individual splice isoform can be found in Supplementary Table 2.

**Supplemental Item 2.**
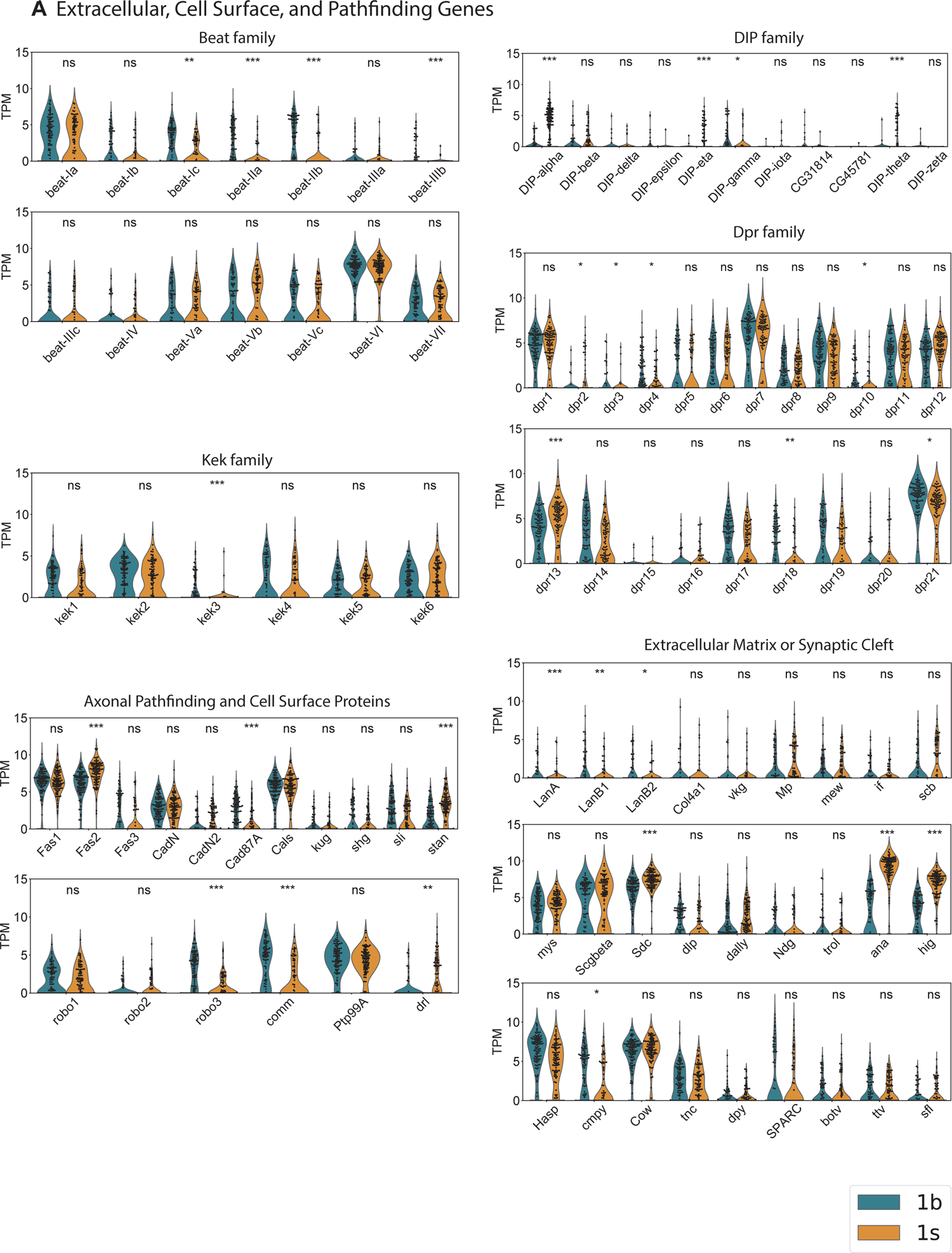
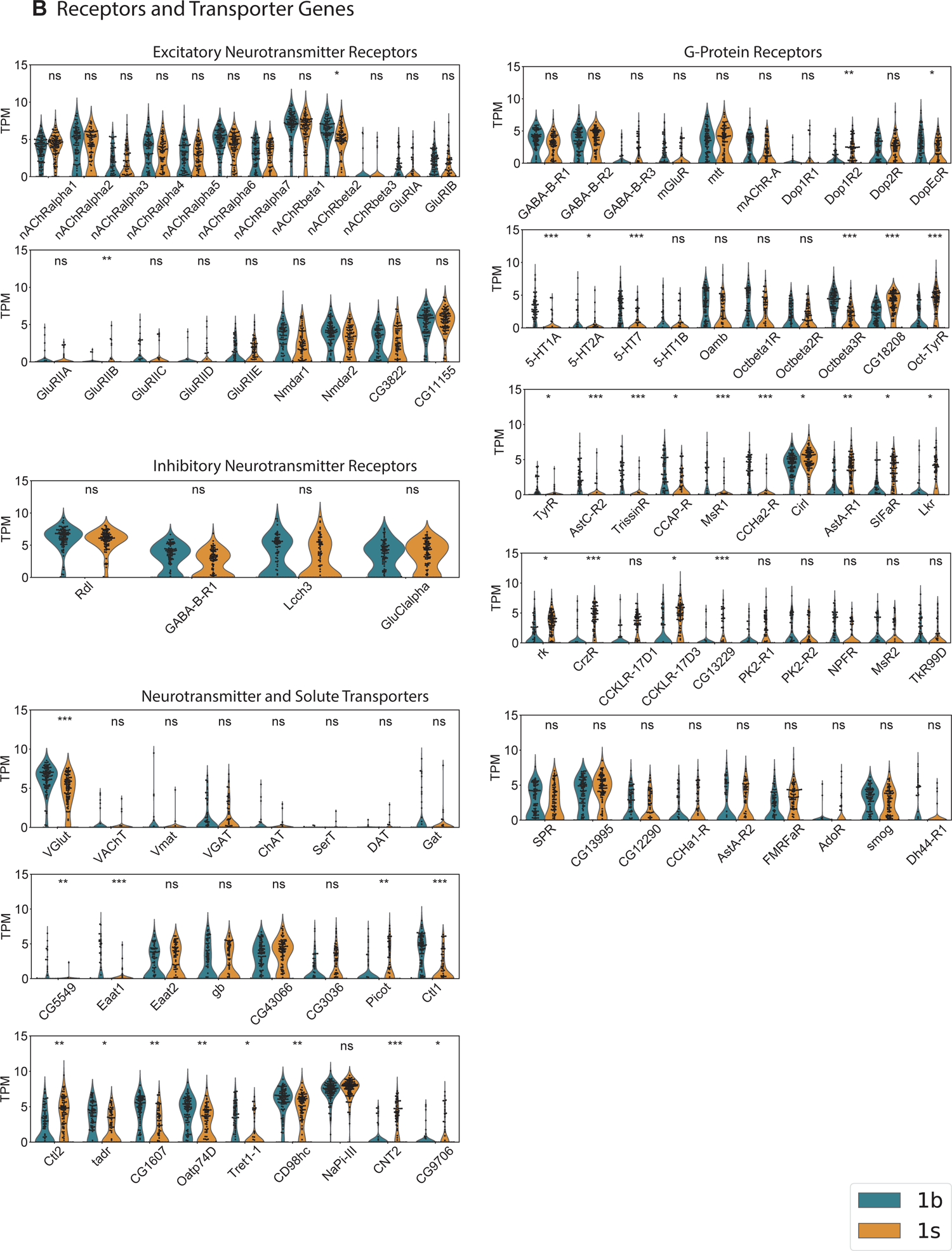
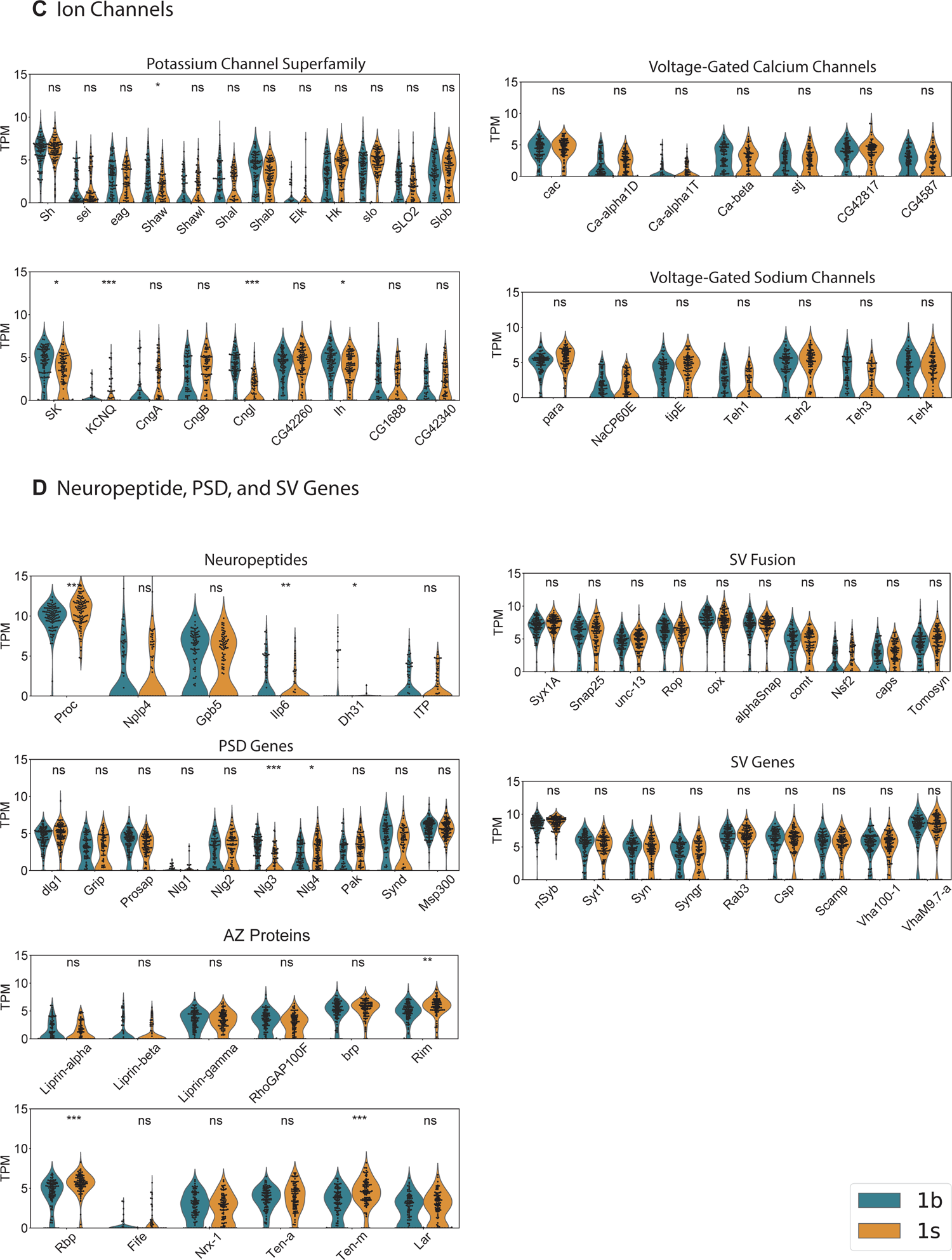
Violin plots of selected DEGs in MN1-Ib and Is. For each gene, expression level (in Transcripts per Million, TPM) of the RNA in each sequenced cell (black dots) is shown. Genes are manually grouped into gene categories for comparison of expression level across member. Violin plots are shown for Ib (blue) and Is (orange) cells. Above each gene is the statistical significance of the Ib versus Is expression level difference (adjusted p-value from the RNA sequencing Wald statistic). Numerical data for each gene can be found in Supplementary Table 1 (all genes) and Supplementary Table 3 (selected genes are sorted into functional categories).

**Supplemental Item 3.**
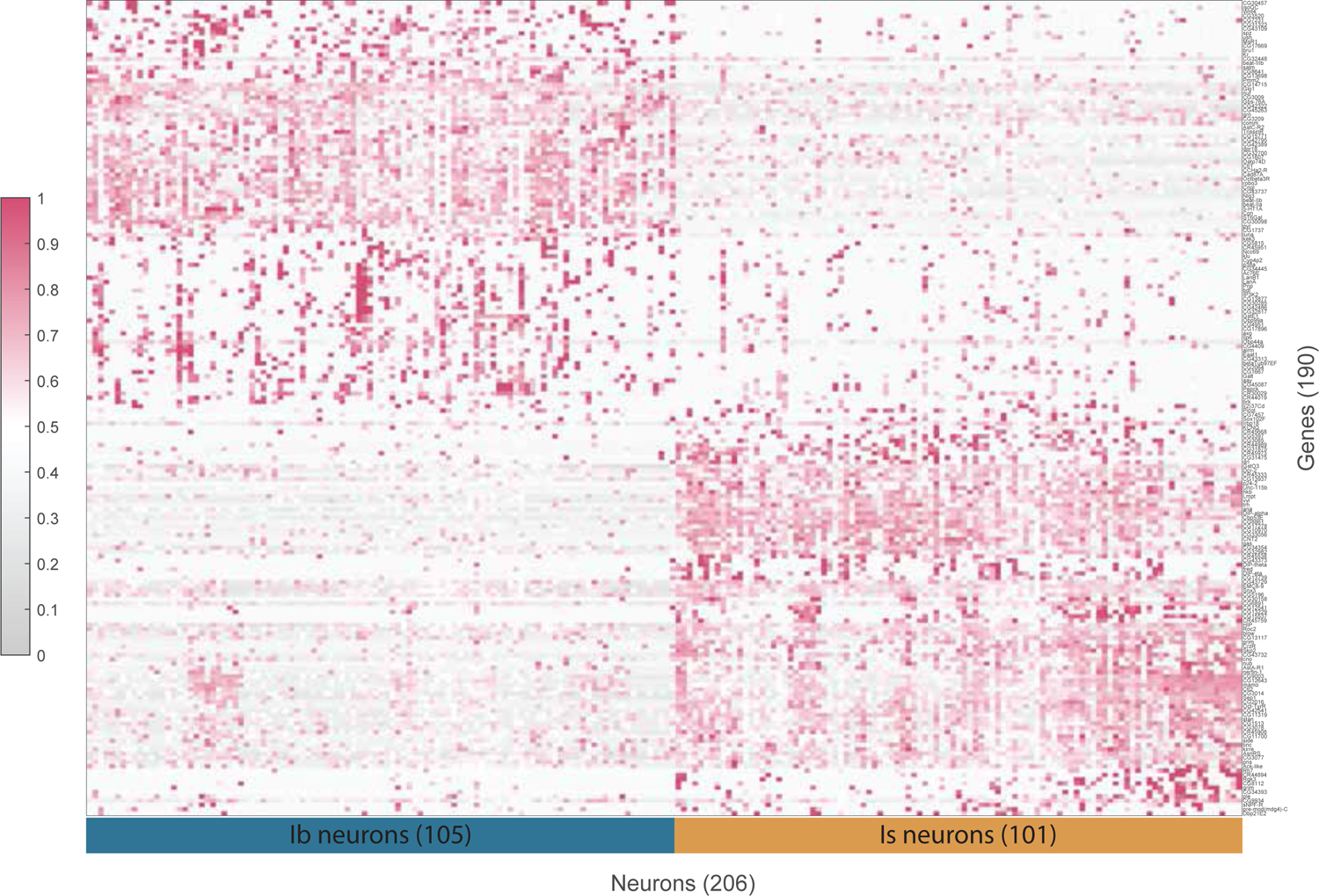
Plot of highly specific MN1-Ib and Is enriched DEGs. Genes which are expressed broadly in one cell type but narrowly in the other are of particular interest given they may encode proteins that drive distinct Ib and Is MN properties. To isolate this group, DEGs were classified as having either “on” (over 70% of Ib or Is MNs express the gene) or “off” (less than 20% of Ib or Is MNs express the gene) expression (“on” or “off” genes analysis adopted from Shrestha et al., 2018, Cell 174, 1229–1246, Figure 1). DEGs that were “on” or “off” in both cells types were not included. Narrowing the group to those with an adjusted RNAseq p-value of less than 0.01 identified 190 DEGs fitting the criteria. These genes are plotted in a clustergram (column clustering method with standardized rows) showing the expression level of each gene (transcripts per million, red shading) for each Ib and Is MN. d it’s difference in expression between Ib and Is cells.

**Supplemental Item 4.**
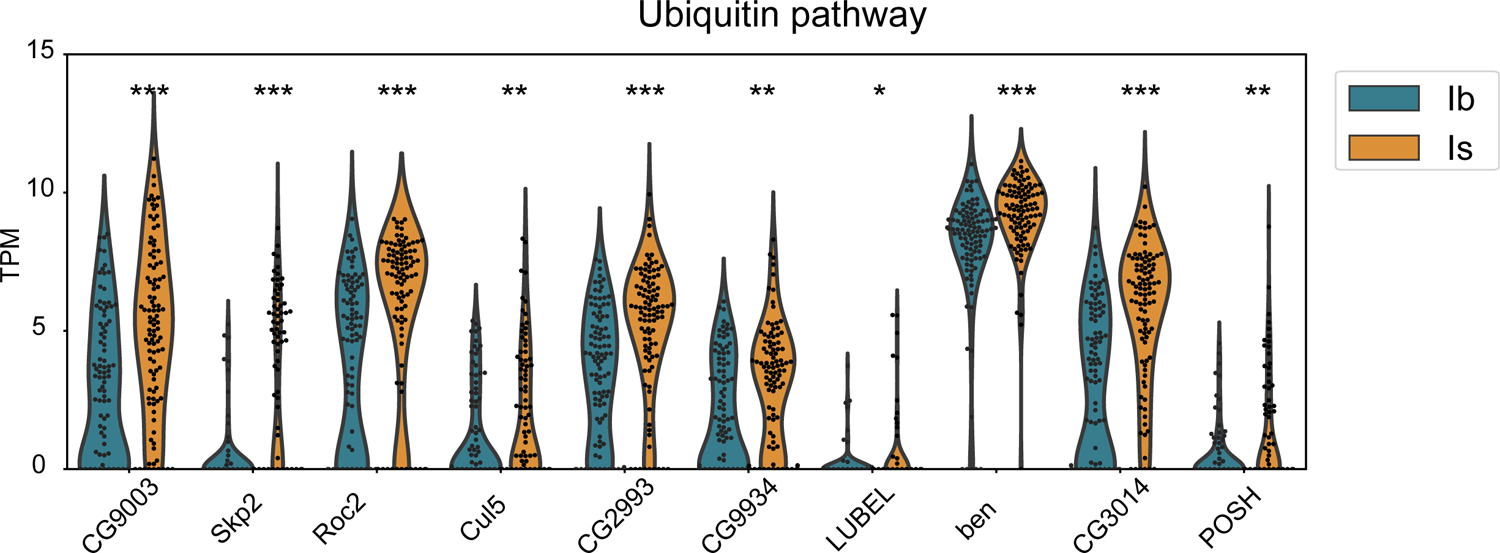
Violin plots of selected DEGs encoding components of the ubiquitin pathway. For each gene, expression level (in Transcripts per Million, TPM) of the RNA in each sequenced cell (black dots) is shown.

**Supplemental Item 5.**
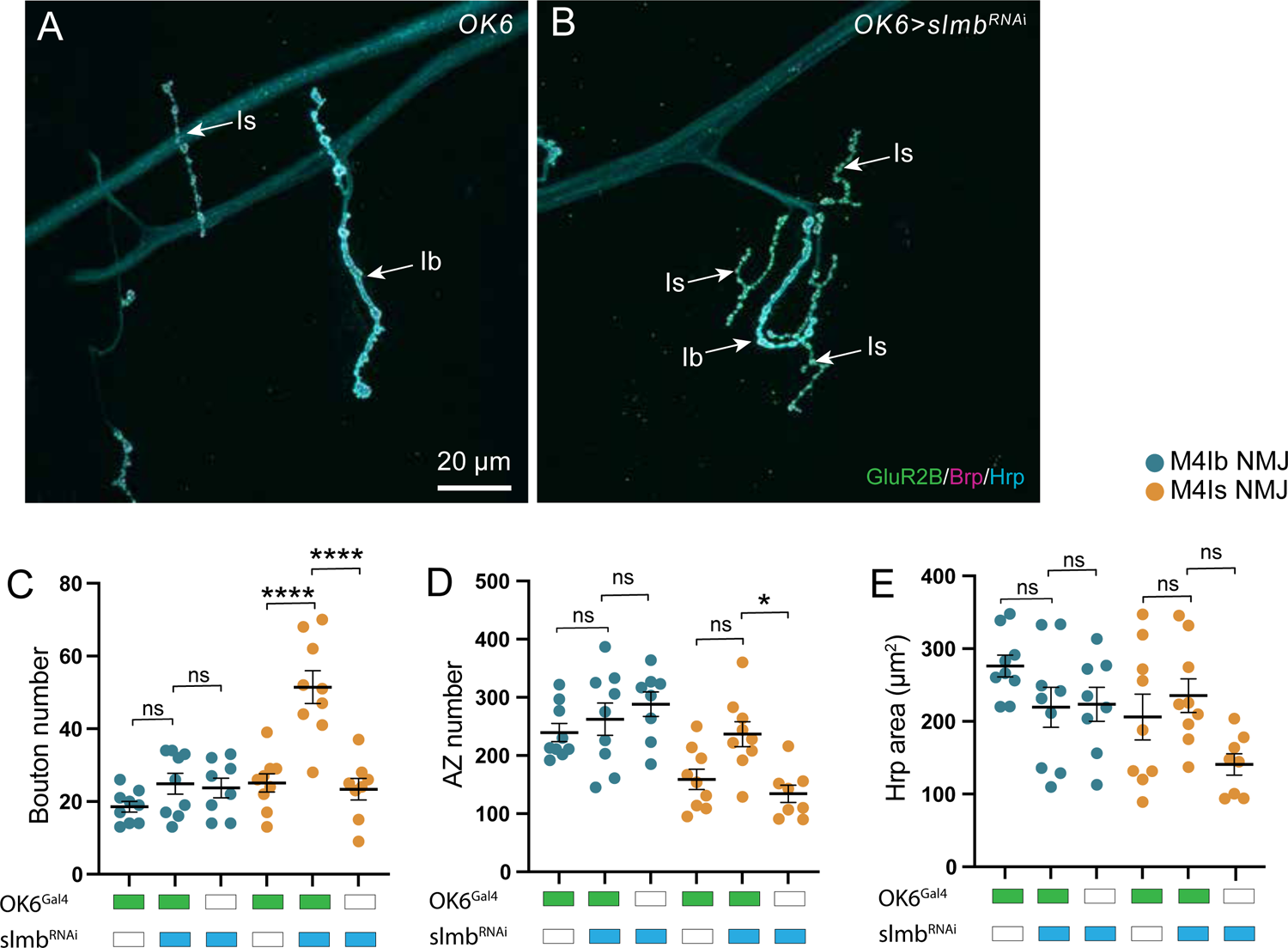
Differential regulation of Ib and Is bouton growth and AZ number in *Slmb* mutants. (A) Representative confocal images of Ib and Is 3^rd^ instar larval muscle 4 NMJs in OK6-Gal4 controls. (B) Representative confocal images of Ib and Is 3^rd^ instar larval muscle 4 NMJs in OK6>slmb^RNAi^. Larvae in panels A and B were immunostained for GluR2B, Brp and Hrp. (C) Quantification of bouton number at 3^rd^ instar larval muscle 4 NMJs (Ib-blue, Is-orange) in controls and OK6>slmb^RNAi^. (D) Quantification of AZ number in controls and OK6>slmb^RNAi^. (E) Quantification of Hrp area (µm^2^) in controls and OK6>slmb^RNAi^. One-way ANOVA with Tukey correction was used to determine significance. \**p* < 0.05, \*\**p* < 0.01, \*\*\**p* < 0.001, \*\*\*\**p* < 0.0001, ns not significant. n=9 NMJs, 5 larvae for panels C-E.

## METHODS

### KEY RESOURCES TABLE

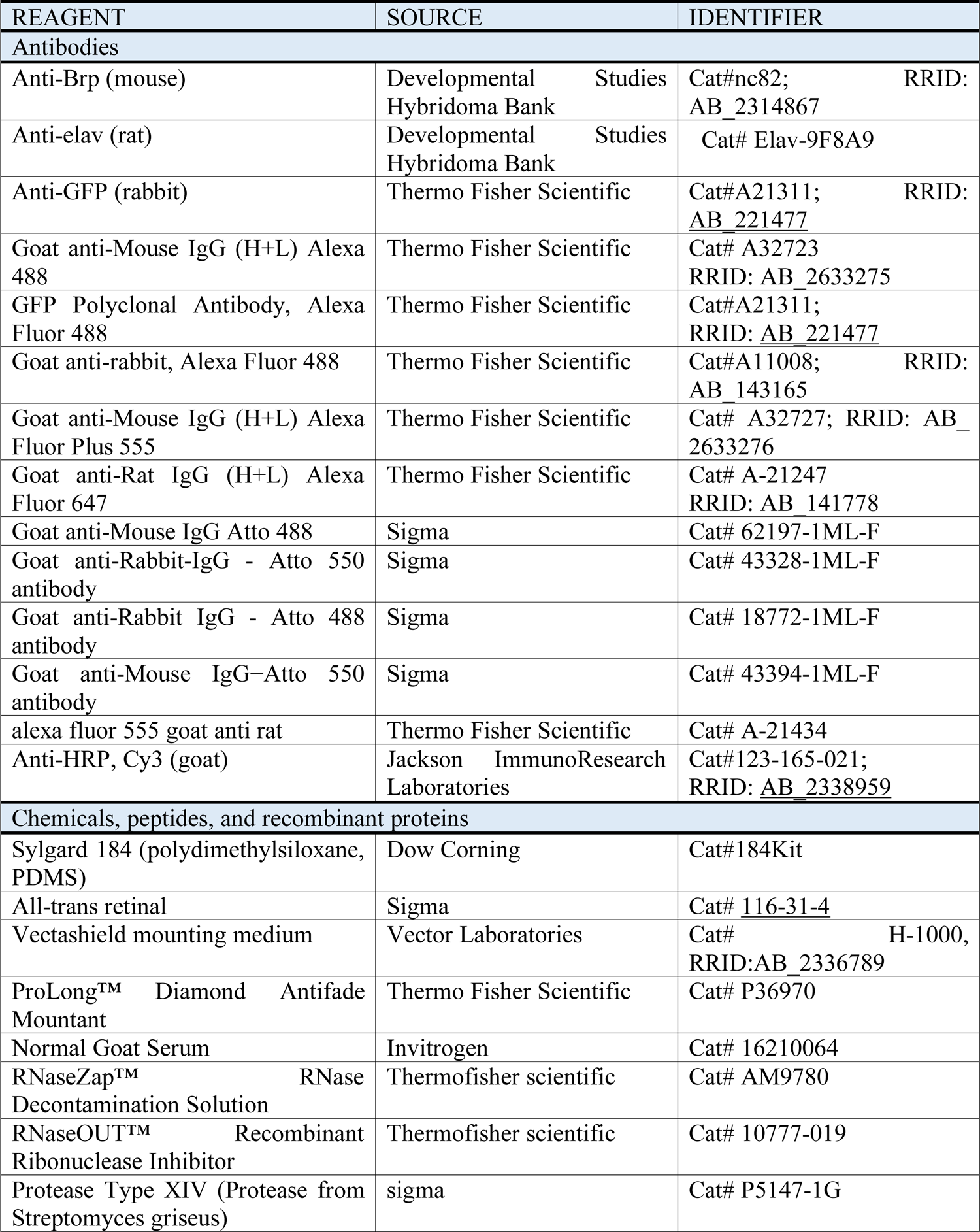
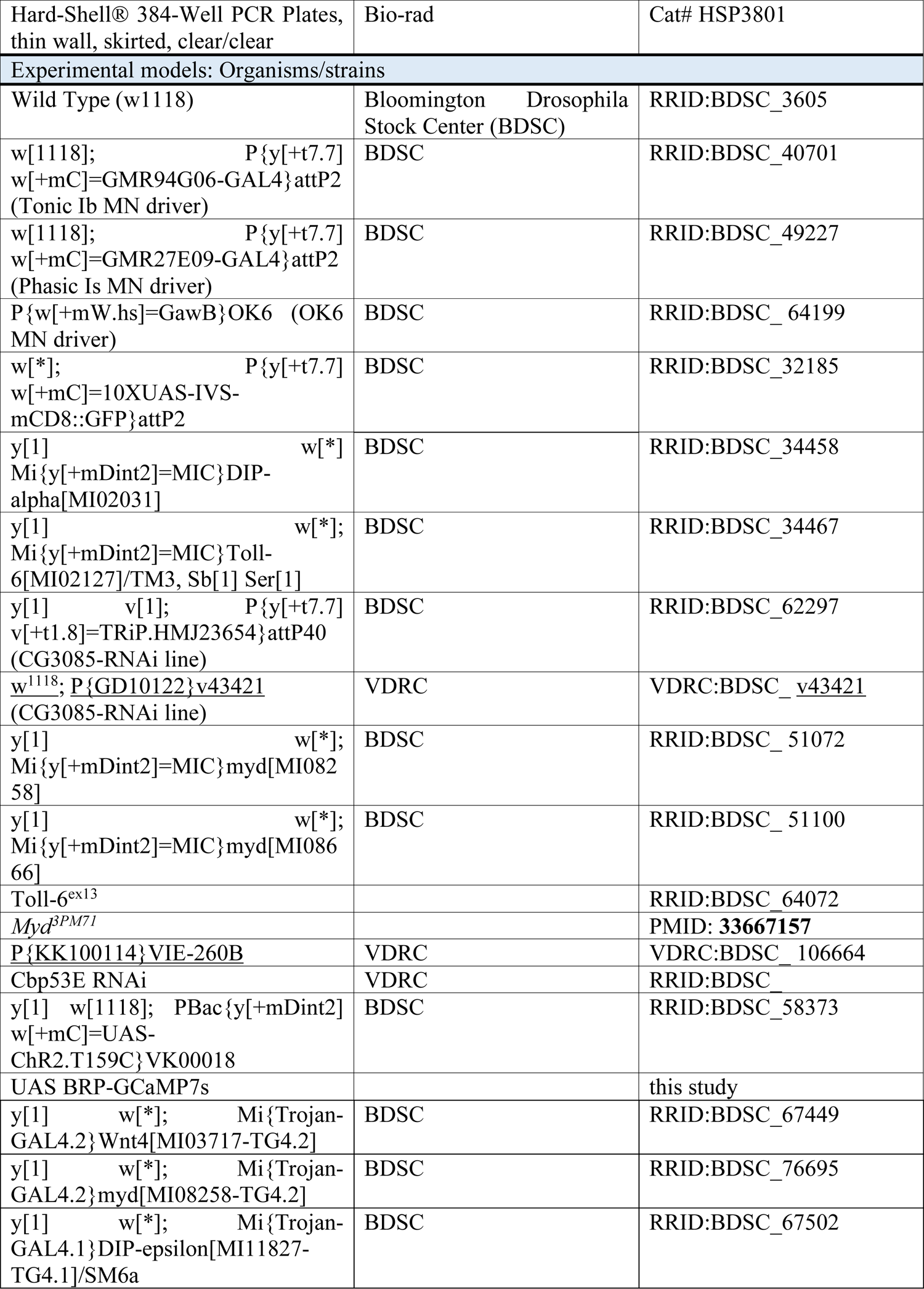
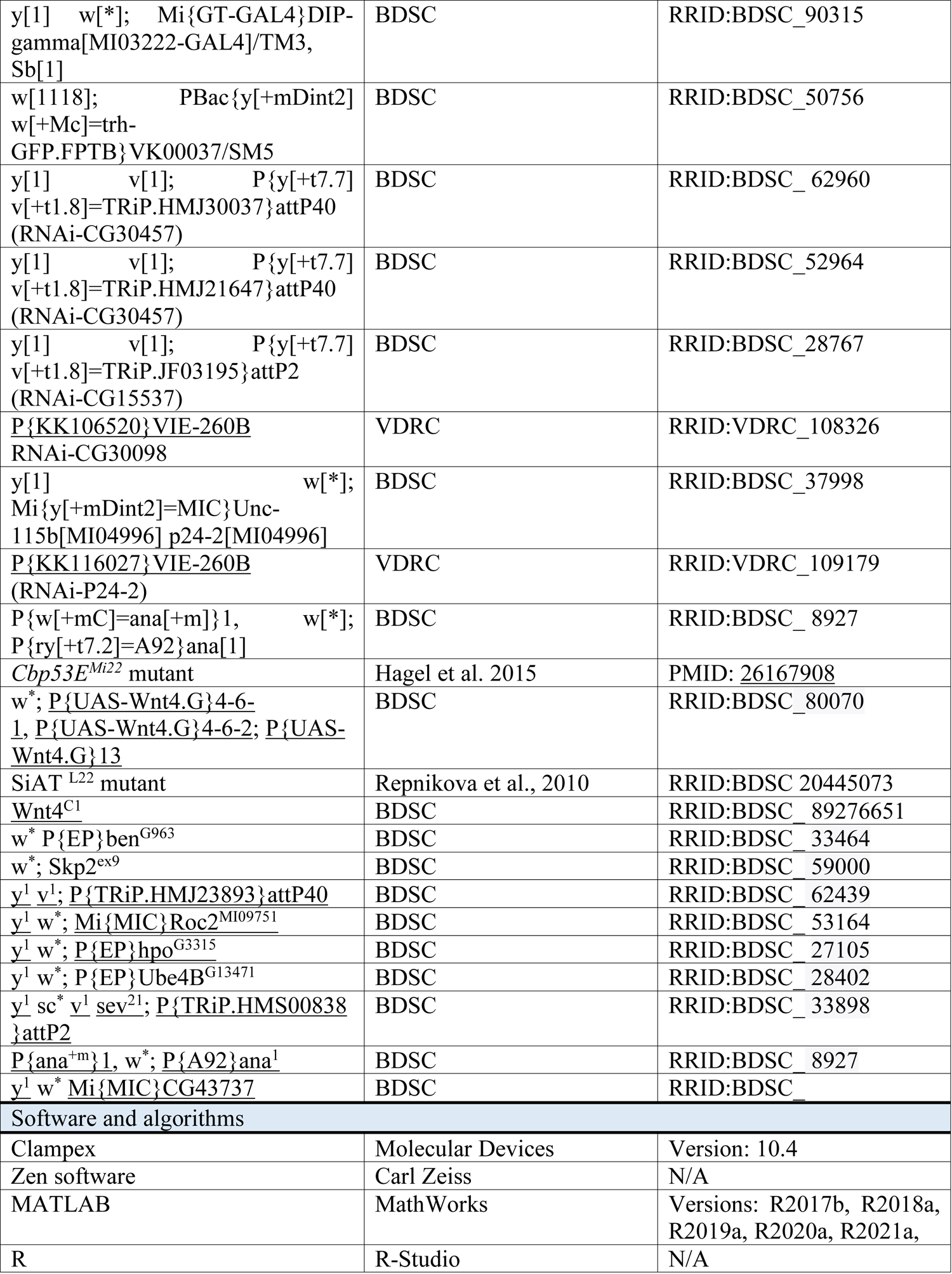
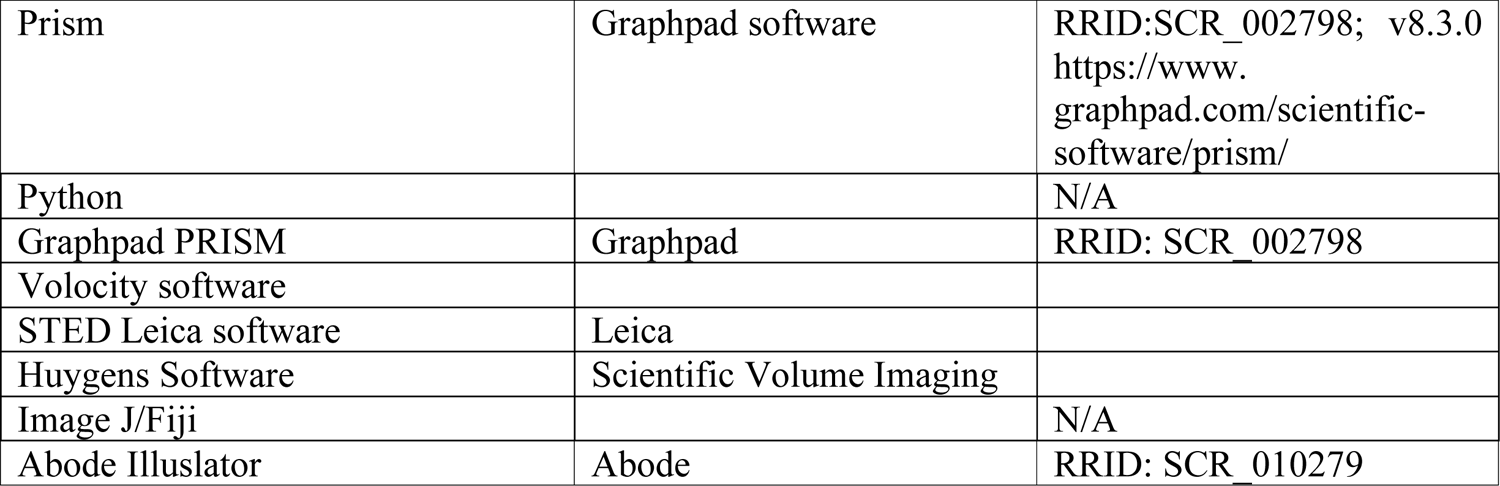

### Experimental model and subject details (*Drosophila* stocks)

Fly stocks were raised at 25°C on standard molasses food in an incubator with 12 hr day/night cycle. Actively crawling 3^rd^ instar larvae that were size and age matched were used for all experiments. *w^1118^* or OK6^Gal4^ driver lines were used as control as indicated. Fly strains used in the study are indicated in the resource table.

### Larval fillet preparation

Wandering 3^rd^ instar larva were dissected in cold Ca^2+^ free hemolymph-like-3 (HL3) saline solution consisting of the following in (mM): 70 NaCl, 5 KCl, 10 NaHCO3, 10 MgCl2, 115 sucrose, 4.2 trehalose, and 5 HEPES (pH 7.2). Evoked synaptic potentials (EJPs) were recorded from muscle 1 using a modified fillet preparation. 3^rd^ instar larvae were pinned down in a sylgard plate and a lateral cut was made from the anterior to the posterior end. CNS, gut, and fat bodies were gently removed. After the dissection, saline was replaced with HL3 containing 0.5 mM calcium prior to electrophysiology.

### Optogenetic stimulation

To stimulate Is or Ib MN NMJs, UAS^ChR2-T159C^ was expressed using Is- or Ib-Gal4 drivers. 3^rd^ instar larvae were grown in standard fly food supplemented with 0.5 mM all-trans-retinal. To prevent photoconversion of all-trans-retinal, aluminum foil was wrapped around food vials and larva were then grown under standard conditions as described. Optogenetic stimulation of Ib or Is MN terminals was carried out using a CoolLED pE-4000 system to deliver short blue light pulses (470 nm). Light pulses were applied via a 10X Olympus water objective for a duration of 50 to 100 microseconds with an interval 5 seconds to elicit unitary synaptic potentials in a reliable manner.

### Sharp electrode current-clamp electrophysiology

Sharp electrodes were backfilled with 3M KCl using gel loading tips. The resistance of the sharp electrodes was in the range of 20 to 30 mOhm after filling with 3M KCl. Optically evoked synaptic potentials were measured in current-clamp mode using a Multiclamp 700b amplifier (Axon instruments), digitized with Digidata 1440 (Axon instruments), and sampled with pClamp10 software. Signals were acquired at 10 kHz and low pass filtered at 2 kHz. Synaptic potentials were recorded from muscle 1 NMJs in abdominal segments 3 and 4. Resting Vm and muscle input resistance were monitored at the beginning and end of each recording. Recordings were excluded if resting Vm and muscle input resistance (Rin) changed by more than 20%.

### Isoform PatchSeq protocol

To prevent contamination of Patch-Seq samples, the fly station, lab bench, dissection microscopes, patch-clamp rig, perfusion apparatus, and microelectrode laser puller were cleaned with 70% ethanol, followed by RNase zap. Borosilicate microcapillaries, glassware, milli-Q water and other supplies were autoclaved to avoid contamination. Single Ib and Is MN or muscle 1 or 4 samples were collected using patch electrodes pulled using autoclaved borosilicate microcapillaries. 3^rd^ instar larval fillets were dissected in ice-cold sterile external saline consisting of (in mM): 103 NaCl, 3 KCl, 5 TES, 8 trehalose, 10 glucose, 26 NaHCO3, 1 NaH2PO4, 4 MgCl2. The osmolarity of the sterile external saline was adjusted to 275-280 mOsm. The sterile saline was bubbled with carbogen gas (95% O2 and 5% CO2) and adjusted to pH 7.3. During PatchSeq sample collection, sterile external saline was continuously superfues over the CNS of intact 3^rd^ instar larvae at a flow rate of 2-3 ml per minute. Following sample collection, 70% ethanol was run through the custom-made perfusion system to maintain sterile conditions and avoid contamination.

To gently disrupt the glia sheath surrounding Ib and Is MNs, 0.5 to 1% protease (Type XIV, Streptomyces griseus, Sigma) was applied using a whole-cell electrode (2 to 3 µm tip size). Glial sheath disruption was carefully observed using microscope optics, followed by protease treatment. After glia sheath disruption, residual protease was quickly washed out by increasing the flow rate of external saline to 5-7 ml per minute for 5-10 minutes, and the flow rate was returned back to 2-3 ml per minute for the remaining sample collection. Residual debris and glial sheath covering MNs were removed using a cleaning electrode (3 to 4 µm tip size) filled with external sterile saline to ensure a clean surface before MN samples were collected.

For PatchSeq collection, sterile whole-cell electrodes were filled with 1 µl of nuclease-free water containing 20 units/ul of RNAse inhibitor. To visualize the cell bodies of Is and Ib MNs, 10XUAS-mCD8-GFP was expressed using Is or Ib Gal4 drivers. MN cell bodies were visualized using a BX51W Olympus microscope with a 40X water objective. Whole-cell electrodes were targeted to the MN cell bodies by applying continuous positive pressure using mouth suction. Somatic and nuclear contents of single MNs were gently collected into the electrode. After sucking up the somatic and nuclear content of single MNs, the electrode holder was carefully removed from the micromanipulator, and the tip of the electrode was broken gently into a 384 microwell in a quadrant pattern. Samples were collected from thoracic (T1-T3) and abdominal segments (A1-A8) from both sexes. To prevent temperature fluctuations, 384 microwell plates were maintained on dry ice in closed thermal coal. MN and muscle samples were collected into a 384 microwell plate in a quadrant fashion, and 96 samples were collected into one plate for library preparation. As a control, 2 or 3 wells were filled with nuclease-free water only. Samples were collected within a time window of 4 to 6 days and stored in a −80 freezer between sample collection. As larval single MN patchSeq was not carried out previously, a pilot RNAseq analysis was performed to determine whether it was feasible to detect RNA profiles from single Ib and Is MNs. The pilot analysis robustly detected gene expression profiles in Is and Ib MNs. Samples from muscle 1 or 4 were collected using low resistance patch electrodes (tip size of 5 to 10 µm) filled with external saline and flash frozen on dry ice together with the MN samples in 384 well plates.

### Single-cell RNA library preparation and Sequencing

RNAseq libraries from single cells were prepared using a modified version of the SMART-seq2 single-cell protocol (Picelli et al., 2013, Trombetta et al., 2014). RNA from single cells was placed in Hard-Shell 384-well plates (Catalog #HSP3801) in minimal volume (<2 uL). cDNA was prepared following a standard protocol using 1/3 recommended volumes for all reagents utilizing a Mosquito HV automated liquid handler (SPT Labtech). Amplified cDNA was purified using 0.75X SPRI beads, quantified using picogreen and spot-checked on a Fragment Analyzer (Agilent). Dual indexed Illumina libraries were prepared from cDNA using a reduced volume NexteraXT reaction (1:12) on the Mosquito HV (Hendricks et al., in preparation). Final libraries were quantified using picogreen and spot-checked on a Fragment Analyzer before pooling and final quantification using qPCR on a Roche LC480II. Libraries were run on an Illumina HiSeq2000 (single end 40nt) or Illumina NextSeq500 (paired-end 75nt).

### Synaptic GCaMP imaging

3^rd^ instar larvae were dissected in Ca^2+^ free HL3. Optical quantal imaging experiments were carried out using HL3.1 containing 20 mm Mg^2+^ and 1.3 mm Ca^2+^ using a confocal microscope (Perkin Elmer) with a spinning-disk confocal head (CSU-X1; Yokogawa), ImagEM X2 EM-CCD camera (Hamamatsu), and 488, 565, and 670 lasers. An Olympus LUMFL N 60X objective with a 1.10 NA was used to acquire GCaMP7s signals from muscle 4 NMJs. To identify individual AZs, a CRISPR Cac-TagRFP (Gratz et al., 2019) strain that endogenously tags the Cac protein was used and membrane tagged (myristoylated) GCaMP7s was expressed in muscles using a Mef2-LexA driver. Prior to quantal imaging, 3D image stacks of muscle 4 NMJs were collected to identify the spatial location of individual AZs denoted by localized Cac-TagRFP fluorescence. To measure evoked release probability of Ib and Is MNs, axons were stimulated using minimal stimulation protocols. Care was taken to recruit both Is and Ib by adjusting the stimulation intensity during imaging experiments. Stimulation was applied using an AMPI Master-8 stimulator at 0.3 Hz frequency for a duration of 200 microseconds for an experimental time window of 5 minutes. Volocity 3D Image Analysis software (PerkinElmer) was used to acquire and analyze data.

### Immunohistochemistry of NMJ and CNS samples

Larval fillets were dissected in ice-cold Ca^2+^ free HL-3 solution as described earlier. The fillet’s CNS was kept intact to validate expression patterns of DEGs at MN somas and NMJs. Larval fillet samples were fixed in 4% formaldehyde for 15-20 minutes at room temperature, washed in PBST (PBS with 0.3% Triton X-100) for 1 hour at room temperature, and incubated in blocking buffer (PBST+ 5% normal goat serum) for 1 hour at room temperature on a shaker. All primary and secondary antibodies were diluted in PBT. Samples were incubated in the primary antibody at 4°C overnight while rocking, with three 20 minutes washes of PBT at room temperature before secondary antibody incubation. Fillet samples without the CNS were incubated in secondary antibody for 2 hours, whereas CNS intact fillet samples were incubated in secondary overnight at 4°C while rocking. Samples were mounted in Vectashield and imaged with a Zeiss confocal microscope using a 60x (NA = 1.3) oil objective. Samples for STED imaging were mounted in prolong diamond mount to improve quality of STED depletion. Primary and secondary antibodies used in this study are listed in the resource table.

### Confocal imaging, image acquisition and processing

A Zeiss LSM 780 laser-scanning confocal microscope (Carl Zeiss) with a 63x/1.3NM oil-immersion objective (Carl Zeiss) was used for data collection for bouton and AZ quantification. Confocal z stacks were obtained in 1 mm intervals at a resolution of 1084×1084. Images were processed with Zen software (Zeiss) to obtain maximum projections. Photoshop (Adobe) or Fiji were used for image rotation and cropping. Adobe Illustrator was used to assemble images and prepare figures.

### STED imaging

Dual or triple channel STED imaging of Ib and Is AZs was carried out using a Leica SP8 confocal/STED 3× microscope (Leica Microsystems, Mannheim, Germany) with 100X, 1.44 NA oil objective and immersion oil (Leica Type F, refractive index 1.5180). For STED, immunohistochemistry protocols were modified to improve nanoscale analysis of AZs. Instead of 20 minutes of 4% PFA fixation, 10 minutes of fixation were used following calibration experiments. After overnight primary antibody incubation, samples were washed 3-4 times in PBST for 1 hour and incubated in Atto-conjugated secondary 488 antibodies, Alexa 555 and Hrp 647. Samples were washed three times for 1 hour in PBST and mounted on slides using prolong diamond mount.

Triple or dual-color sequential STED scans were acquired using the STED microscope with gated detectors and 40 nm x-y resolution. Based on pilot studies, a combination of Atto 488 and Alexa Fluor 565 were selected as optimal for analyzing nanoscale AZ organization as the two dyes were photostable and showed no shifts in excitation of wavelengths, and photobleaching of signals was significantly reduced compared to other tested dye combinations. Images were scanned at a pixel density of 1024 × 1024 (9 nm/pixel). Hrp 647, Alexa Fluor 555, and Atto 488 were excited with 633 nm, 555 nm, and 488 nm white light lasers, respectively, in the same sequential order at 2-5% 1.5 mW laser power. We depleted Atto 488 and Alexa Fluor 565 signals using 592 nm (75% of maximum power) and 660 nm (25% of maximum power) time-gated depletion lasers. During STED scanning, line accumulation of four times and frame averaging of three times was used. Eight times zoom was used to compare the nanoscale organization of AZs and other key synaptic proteins.

### TEM sample preparation and imaging

TEM sample preparation was carried out as described previously (Guan et al., 2020). 3^rd^ instar larval preparations were dissected in Ca^2+^-free HL3.1 solution and fixed in freshly prepared fixative containing 1% glutaraldehyde, 4% formaldehyde, and 0.1 m sodium cacodylate for 10 min at room temperature. Fixed samples were microwaved in a BioWave Pro Pelco (Ted Pella, Inc, Redding, CA, USA) in the following order: (1) 100W 1 min, (2) 1 min off, (3) 100W 1 min, (4) 300W 20 secs, (5) 20 secs off, (6) 300W 20 secs, with steps 4-6 repeated twice. Following microwaved steps, samples were incubated for 30 min at room temperature in fixative. Samples were washed in 0.1 M sodium cacodylate and 0.1 M sucrose and were stained for 30 min in 1% osmium tetroxide and 1.5% potassium ferrocyanide in 0.1 M sodium cacodylate solution. Samples were washed in 0.1 M sodium cacodylate and stained for 30 mins in 2% uranyl acetate. Samples were then embedded in epoxy resin (Embed 812; Electron Microscopy Sciences) and dehydrated through a graded series of ethanol and acetone. 50–60 nm range thin sections were collected on Formvar/carbon-coated copper slot grids and contrasted using lead citrate. A Tecnai G2 electron microscope (FEI, Hillsboro, OR, USA) equipped with a charge-coupled device camera (Advanced Microscopy Techniques, Woburn, MA, USA) was used to carry out TEM at 49,000 × magnification at 80 kV.

Ib and Is boutons at muscle 6/7 NMJs were analyzed to compare Ib and Is synaptic features. Ib and Is innervation were identified by there position on muscle 6/7 and associated SSR size. Only micrographs with bouton diameters larger than 2 µm were included in analysis for 1b boutons. For SV counting, T-bars at Ib and Is boutons were identified and Photoshop was used to draw concentric circles with 100 nm and 200 nm radii with the T-bar at the center. To quantify vesicle density, Volocity software was used to measure the area of the bouton and quantify the total number of SVs within it.

### Data exclusion criteria

For synaptic GCaMP imaging experiments, data was excluded if larvae underwent significant muscle contractions. For electrophysiology experiments, data was excluded based on resting Vm (if resting Vm is above −60 vm) and input resistance threshold. In some cases, data with a sudden drop in resting Vm and Rin were also excluded from the analysis, as were samples where less than 20 traces were obtained during the recording. Patch-Seq samples were excluded with low read number when less than 3000 genes were detected based on alignment to the *Drosophila* genome.

### Replication of results

All experiments were carried out in at least three biological replicates, and all crosses were set up at least twice to obtain reproducible results from replicate to replicate. STED imaging was repeated several times to test variables such as duration of PFA fixation, mounting conditions, intensities for depletion laser, and concentrations of immunostaining reagents to exclude the possibility that differential nanoscale organization of Ib and Is AZs was due to any specific imaging condition. Differences Ib and Is nanoscale organization was highly reproducible in all biological replicates.

## QUANTIFICATION AND STATISTICAL ANALYSIS

### Electrophysiology data analysis

All electrophysiological recordings were analyzed using clampex software and custom-written MATLAB codes. To calculate the average amplitude of evoked EJPs, at least 20 evoked traces were acquired at 0.2 Hz and the average amplitude was determined using Clampex software. To exclude the possibility that optogenetically evoked response were graded responses, simultaneous electric and optogenetic stimulation were alternated, and pharmacology was used to confirm optogenetic responses were absent following action potential blockage.

### Synaptic GCaMP imaging data analysis

Analysis was performed as described in (Akbergenova et al., 2018). Briefly, the position of Cac-TagRFP puncta was detected and uniform ROIs were assigned to each puncta. myrGCaMP7s flashes were assigned to ROIs by proximity. Only events occurring within 0.8 µM distance from ROI centers were analyzed. Experiments with a significant amount of image movement were not included in our analysis.

### STED image analysis

All STED images were deconvolved using the deconvolution module in the Huygens software by applying the classical maximum likelihood estimation (CMLE) deconvolution algorithm to increase signal-to-noise. As Huygens software uses a signal reassignment-based deconvolution algorithm, it is feasible to carry out intensity measurements and comparisons between samples that were collected under the same imaging conditions. Deconvolved images were analyzed in ImageJ (Fiji) software to calculate the area of Is and Ib AZs. To determine the geometric organization of Is and Ib AZs, Wayne Rasband’s ImageJ circularity plugin was used, where circularity is measured as *circularity = 4pi (area/perimeter^2)*. If the CI index is 1, AZs are perfectly circular, and if CI index is 0 or close to 0, AZs are non-circular. CI index analysis robustly classified the geometric organization of Ib and Is AZs. ROIs were manually drawn around AZs to calculate area, perimeter, circularity index. To show differences in nanoscopic organization of AZs, intensity profile analysis, a complementary approach to CI index analysis, was performed. Intensity profiles of Ib and Is AZs were are also different, supporting distinct Ib and Is AZ structure. AZs lacking a core and lateralized AZs were excluded from the analysis.

### Dataset Processing and Single-Sample Gene Set Enrichment Analysis

Paired-end reads were aligned to the *Drosophila melanogaster* transcriptome derived from assembly BDGP6 with Ensembl annotation release 92 and STAR version 2.5.3a (Dobin et al., 2013). Gene expression was quantified with RSEM version 1.3.0 (Li and Dewey 2011) with the calc-pme option. Raw transcript per million (TPM) values were offset by +1 and transformed to log 2 space and hierarchical clustering was done with protein coding genes and manual examination of the results was used to exclude outlier cells from downstream analysis. Integer count data was prepared for genes and isoforms from the posterior mean counts results and the data were used as input for differential expression testing and Seurat-based analyses.

### Differential Gene Expression Analysis

Integer count values derived from RSEM processing were used as input to differential expression analysis with DESeq2 (version 1.24.0, Love et al., 2014) using normal log fold change shrinkage. Differentially expressed genes or isoforms were defined as those having an absolute log2 fold change >= 1 and an adjusted p value <= 0.05. Differential expression data was visualized using TIBCO Spotfire Analyst version 7.11.1.

### Visualization and Clustering of Single Cell Data

Integer count data was imported into Seurat (version 3.0.0, Stuart et al., 2019) running under R version 3.6.0 with and dimensionality reduction plots were prepared with umap (version 0.2.2.0) using dims of 1:20. Additional R packages include cowplot (version 0.9.4), ggplot2 (version 3.1.1) and dplyr (version 0.8.0.1).

### Gene ontology analysis

To determine if sets of related genes or genes with related functions were differentially regulated in 1b and 1s MNs, a list of all genes rank-ordered by RSEM statistic was uploaded to the STRING database (https://string-db.org/). The search tool for Functional Enrichment Analysis was used with the organism set to *Drosophila melanogaster* with a 5% false discovery rate.

### RNAseq analysis, heatmaps, and violin plots

For visualization, Matlab was used to plot heatmaps and scatterplots. The Seaborn statistical data visualization package (https://seaborn.pydata.org/) was used to make combined violin plots and swarm plots of transcript level data, running in the Spyder 4.2.1 IDE using Python 3.7.

### Statistical analysis

Data represents mean ± SEM. One-way ANOVA with Tukey correction was used to determine significance for multiple comparisons. Student t-test was used for two bar graphs. \**p* < 0.05, \*\**p* < 0.01, \*\*\**p* < 0.001, \*\*\*\**p* < 0.0001, ns not significant. All reported values are mean ± SEM.

### RNAseq data availability

All Isoform-Patchseq RNA profiling raw data is available at NCBI under the GEO accession # GSE222976.

## Notes

### Competing Interest Statement

The authors have declared no competing interest.

